# TERT drives liver tumorigenesis beyond telomere elongation

**DOI:** 10.64898/2026.02.02.703327

**Authors:** Laura Braud, Julien Vernerey, Arnaud Guille, Pierre Cordier, Clémence Ginet, Tom Egger, Manuel Bernabe, Dmitri Churikov, Quentin Da Costa, Aïda Meghraoui, Chantal Desdouets, Li Gu, François Bertucci, Christophe Lachaud, Vincent Géli

## Abstract

We generated two mouse models, p21^⁺/Tert^ and p21^⁺/TertCi^, expressing either telomerase reverse transcriptase (TERT) or a catalytically inactive variant under the control of the p21 promoter. By 18–20 months of age, approximately 25% of mice from both genotypes developed liver tumors with histopathological features resembling human hepatocellular carcinoma (HCC). Whole-exome sequencing identified activating Ctnnb1 mutations and recurrent PP1 subunit alterations in p21^⁺/Tert^ tumors, whereas p21^⁺/TertCi^ tumors harbored activating Hras^Gln61Lys^ mutations associated with elevated C>A transversions. Both models exhibited chromosomal aberrations commonly observed in human HCC. Transcriptomic analyses revealed that β-catenin–activated tumors recapitulated gene expression signatures of human HCC, while MAPK-mutated tumors showed profiles consistent with MAPK/ERK pathway activation. Metabolically, both genotypes demonstrated increased glycolysis and suppression of gluconeogenesis, including downregulation of FBP1, but expressed distinct NRF2 target genes. Spatial profiling further revealed reduced HNF4α-positive hepatocytes across tumors, independent of Hnf4α transcription, and markedly diminished immune cell infiltration particularly in β-catenin–activated tumors. Collectively, these findings uncover telomere-independent functions of TERT and identify molecular and metabolic features with potential relevance for predicting immunotherapy response.

## Introduction

Hepatocellular carcinoma (HCC) is the third leading cause of cancer-related mortality worldwide. It arises in the setting of chronic liver diseases such as chronic hepatitis B or C infection, cirrhosis, and metabolic syndrome which are frequently accompanied by persistent inflammation (Zucman-Rossi *et al*, 2015; Nault *et al*, 2019). Within this pathological environment, the accumulation of driver mutations leads to the disruption of key signaling pathways, ultimately driving hepatocarcinogenesis (Schulze *et al*, 2015). Among the most frequently deregulated are the Wnt/beta-catenin and p53 pathways (Rebouissou *et al*, 2016; Fujimoto *et al*, 2015). In addition, multiple other pathways contribute to HCC development, including recurrent amplifications of MYC and cyclin-dependent kinase genes, and mutations that activate the Sonic Hedgehog, RAS/MAPK, PI3K-Akt, and oxidative stress response pathways (Chen *et al*, 2010; Delire & Stärkel, 2015; Zhang *et al*, 2016).

Frequent mutations in hepatocellular carcinoma (HCC) occur early in carcinogenesis within the promoter region of the TERT gene, which encodes the catalytic subunit of telomerase (Nault *et al*, 2013, 2014). Various genetic and epigenetic mechanisms, including TERT promoter mutations, hepatitis B virus integration, chromosomal rearrangements, and promoter methylation changes, lead to aberrant upregulation of TERT in the majority of HCC cases (Totoki *et al*, 2014). Traditionally, telomerase reactivation in HCC is considered as a compensatory response to critical telomere shortening preventing senescence and apoptosis especially in cells harboring p53 mutations (Farazi *et al*, 2006). However, emerging evidence indicates that TERT also performs non-canonical functions beyond telomere elongation that may promote tumorigenesis (Ségal-Bendirdjian & Geli, 2019). Depending on cellular context, TERT has been implicated in the regulation of major oncogenic pathways, including WNT/beta-catenin, NF-κB, and MYC (Park *et al*, 2009; Ghosh *et al*, 2012; Koh *et al*, 2015). Recent studies in murine models of chronic liver inflammation have shown that TERT can enhance NF-κB promoter activity and drive liver tumor formation, particularly in the absence of functional p53 (Mishima *et al*, 2024). Despite these findings, the non-canonical roles of TERT in HCC remain poorly understood.

We developed two related mouse models, p21^⁺/Tert^ and p21^⁺/TertCi^, in which either wild-type TERT or a catalytically inactive TERT mutant (TERT^Ci^) is expressed under the control of the *Cdkn1a* (p21) promoter. This strategy was designed to establish a tightly regulated feedback loop, enabling telomerase expression specifically in p21-positive cells. Unexpectedly, we found that both TERT and TERTCi suppress p21 expression and attenuate cellular senescence either in the lungs of aged mice (Lipskaia *et al*, 2024) or in the adipose tissue of young obese mice fed a high-fat diet (Braud *et al*, 2025). These results suggest that TERT can inhibit senescence through mechanisms independent of its canonical telomere-lengthening activity. Overall, the p21^⁺/Tert^ and p21^⁺/TertCi^ models promote TERT expression in pre-senescent cells and facilitate senescence bypass, an early event frequently associated with tumor initiation, particularly in the liver. In this study, we demonstrate that 15% of p21^⁺/Tert^ and p21^⁺/TertCi^ mice develop liver tumors by 18 months of age, exhibiting characteristics closely resembling those of human hepatocellular carcinoma (HCC). Through these two mouse models, we identify a telomere-independent role for TERT in liver tumorigenesis and immune modulation. These findings reveal non-canonical TERT functions and highlight molecular and metabolic biomarkers relevant to HCC pathogenesis and immunotherapy response.

## Results

### p21^+/Tert^ and p21^+/TertCi^ mice develop hepatocellular carcinomas with features of human HCC

We found that by 18–20 months of age, both p21^⁺/Tert^ and p21^⁺/TertCi^ mice predominantly developed liver tumors. We were surprised to find that mice expressing the catalytically inactive TERT^Ci^ developed liver tumors with similar frequency to those expressing active telomerase. The majority of these tumors were of hepatocellular origin and identified as hepatocellular carcinomas (HCC) or hepatocellular adenomas (HCA). A number of hematological malignancies and liver hemangiosarcomas were also observed (Table 1, Figure 1).

**Figure 1.**
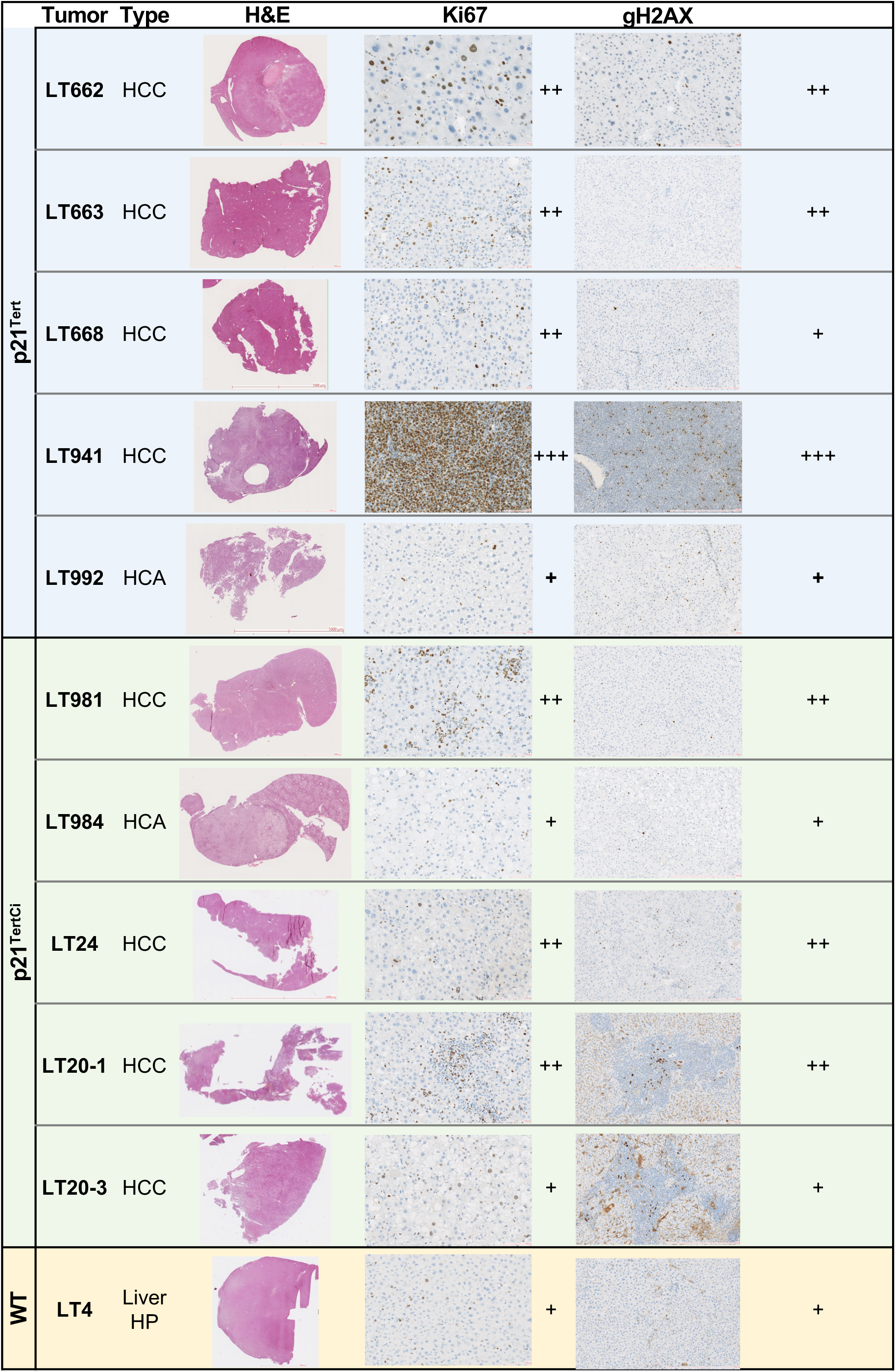
Liver tumors collected from p21^+/Tert^ and p21^+/TertCi^ mice. The histological characteristics of these tumors are summarized in Table 1. Tumor sections were stained with anti-Ki67 and anti-gH2AX antibodies. LT4 represents a case of nodular hepatocellular hyperplasia obtained from an aged wild-type (WT) mouse.

**Table 1.**
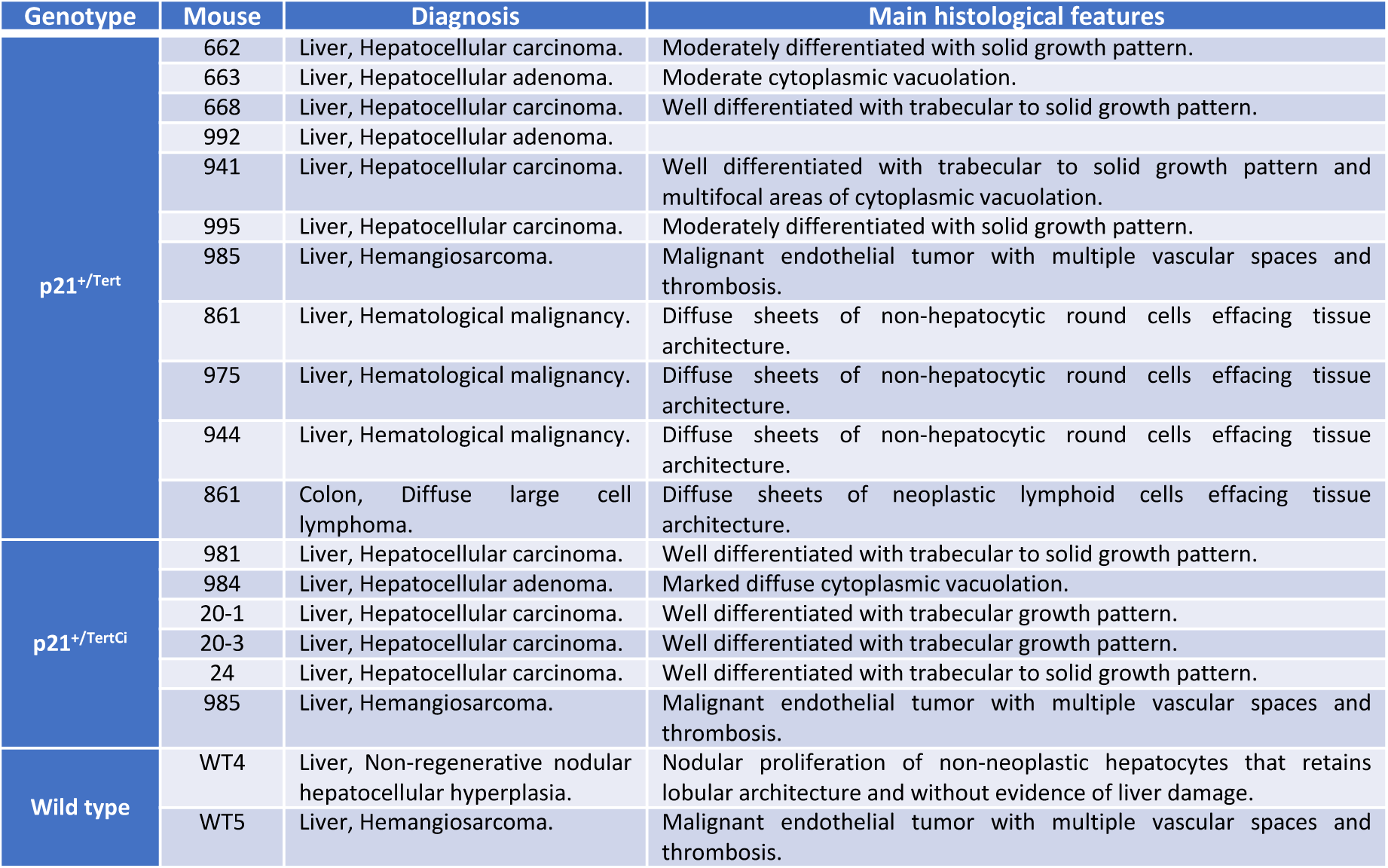
Histological characterization of tumors.

We focused our analysis on hepatocellular tumors, given their predominance. HCCs in p21^⁺/Tert^ and p21^⁺/TertCi^ mice exhibited histomorphological features reminiscent at least to some extent of human HCCs. In general, we noted in most liver tumors, as it is the case for human HCC, a nodular to multi-nodular hepatocellular neoplastic lesion with loss of lobular architecture leading to destruction and rarefaction of adjacent parenchyma, although overt tumor invasion was more difficult to identify (Table 1). The murine HCCs were not encapsulated. We found that the growth pattern of HCCs in the transgenic mice was most often ’micro-*trabecular*’ (neoplastic hepatocytes forming multiple-cell-thick trabeculae alternating with sinusoidal capillaries), or ’*solid*’ (continuous range of neoplastic hepatocytes with very little tumor stroma), with a frequent combination of both patterns within the same tumor (Table 1). These observations are in agreement with the histopathological aspects of human HCC, in which both growth patterns can be found alone or in association. Likewise, the liver tumors from p21^+/Tert^ and p21^+/TertCi^ mice showed multifocal areas of necrosis, hemorrhage or angiectasia. Of note, other tumor growth patterns described in humans (pseudoglandular, macrotrabecular, scirrhous, etc.) were not observed in the p21^+/Tert^ and p21^+/TertCi^ models. Tumor hepatocytes in both mouse models remained well-differentiated (mimicking mature hepatocytes) with a central round euchromatic nucleus and moderately abundant to markedly abundant granular eosinophilic cytoplasm (Table 1), thus resembling low-grade human HCCs. Accordingly, cellular atypia was mostly mild to moderate with increased nuclear–cytoplasmic ratio, anisokaryosis and occasional karyomegaly and nuclear pleomorphism. In some tumors, neoplastic hepatocytes showed prominent cytoplasmic vacuolation. Overall, the malignant hepatocellular tumors developed in p21^+/Tert^ and p21^+/TertCi^ mouse models were well-to-moderately differentiated hepatocellular carcinomas with trabecular and/or solid growth patterns. Finally, in humans, the adjacent non-tumor liver will be cirrhotic in 80-90% of HCC cases. In our mouse models, hepatic fibrosis of the adjacent parenchyma was minimal to mild, indicating a notable difference from the human disease and its pathogenesis. It is worth noting that hepatocellular hyperplasia was observed in the liver of a wild-type mouse (designated WT4), and was characterized alongside the liver tumors found in p21^⁺/Tert^ and p21^⁺/TertCi^ mice (Figure 1).

To assess tumor cell proliferation, we examined Ki67 expression, a well-established proliferation marker. In human HCC, Ki67 levels are associated with aggressive highly proliferative tumors (Ramos-Santillan *et al*, 2024). Consistent with this, all tumors in our study showed markedly increased Ki67 staining compared to normal liver tissue (Figure 1; (Schaeffer *et al*, 2023)). LT941 exhibited strong Ki67 labeling, reflecting a highly proliferative phenotype whereas LT992 and LT984 both classified as HCA showed minimal Ki67 labeling (Figure 1). All remaining tumors exhibited high Ki67 levels. We also assessed the levels of gH2AX, a marker of DNA double-strand breaks (DSBs) and genomic instability. Elevated DSBs have been reported in neoplastic lesions of HCC (Xiao *et al*, 2014). We observed high gH2AX staining in LT662 and LT941, and in both tumors the extent of 53BP1 labeling paralleled Ki67 expression (Figure 1). Elevated gH2AX levels were also detected in LT20-1 and LT20-3, two HCC tumors arising in the same mouse (see below).

Together, our results show that aged p21^⁺/Tert^ and p21^⁺/TertCi^ mice (>18 months) develop well-differentiated liver tumors, predominantly hepatocellular carcinoma (HCC). The ability of both active and inactive TERT, when expressed under the *p21* promoter, to promote tumor formation suggests that TERT contributes to hepatocarcinogenesis also through telomere independent mechanisms.

### Distinct somatic mutation profiles in p21^+/Tert^ and p21^+/TertCi^ tumors

We characterized the mutational landscape of liver tumors from aged p21^+/Tert^ and p21^+/TertCi^ mice using Whole Exome Sequencing (WES). This analysis identified coding mutations in driver genes and copy number alteration (CNA) (Table S1 and S2). We examined somatic mutations associated with hepatocellular carcinoma (HCC) in both humans and mice (**Erreur ! Source du renvoi introuvable.**A, Table S1, dataset1) (Kan *et al*, 2013; Zhang *et al*, 2014; Totoki *et al*, 2014; Letouzé *et al*, 2017; Connor *et al*, 2018; Chen *et al*, 2024; Lee, 2015). In liver tumors from aged p21^⁺/Tert^ mice, LT662 and LT663 carried activating mutations in *Ctnnb1*, which encodes β-catenin (*Ctnnb1^Asp32Tyr^*and *Ctnnb1^Thr41Ile^*, respectively) (Figure 2A). LT662 also harbored mutations in *Col11a1* (*Glu1155Lys*), present in ∼6.8% of human HCCs (Kan et al., 2013), and in the histone demethylase *Kdm4d* (*Gln73His*). LT663 carried mutations in *Kdm3c* (*Arg278Gly*), a demethylase regulating adipogenic transcription factors (Viscarra et al., 2020), and *Ddias* (*Pro706Leu*), a gene linked to DNA damage-induced apoptosis and elevated in HCC (Im *et al*, 2023). Tumor LT992 showed few mutations and no CNAs but contained two frameshift mutations: one in *Ppp1r9a* (*Leu1273Thrfs*), a regulatory subunit of protein phosphatase 1 (PP1), and another in *PtprE* (Protein tyrosine phosphatase receptor type E). LT992 also had a truncating mutation in *Sucla2* (*Gln87del*), a TCA cycle enzyme. LT941 harbored a missense mutation in the Furin-like domain of *Egfr* (*Egfr^Ser303Tyr^*), a region known to harbor MAPK-activating mutations. Mutations at nearby codon 254 in this domain were frequently found in a carcinogen-induced mouse model of liver cancer (Connor *et al*, 2018). It also carried a splice-site mutation in *Ppp1r1a* (another PP1 subunit) and a *Ptpn5* (*Arg246Gly*) mutation, encoding PP5 (Protein Phosphatase 5) (Figure 2A).

**Figure 2.**
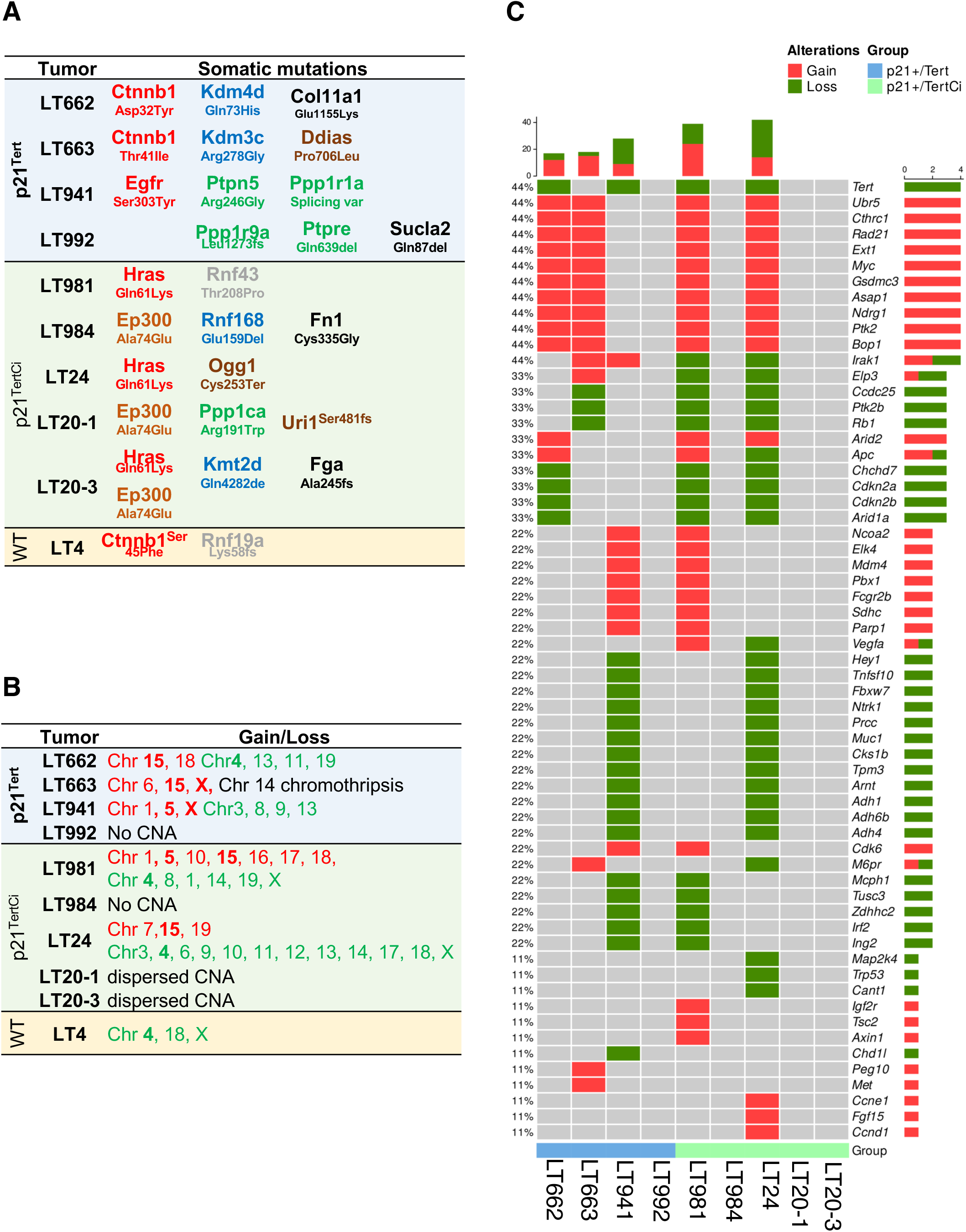
Mutational landscape of liver tumors from aged p21^+/Tert^ and p21^+/TertCi^ mice. **a.** Somatic mutations in genes found altered in HCC at low or high frequency are idincated. All mutations detected in each tumor are listed in Table S1. **b.** Copy number alterations in the liver tumors. Chromosomes with gains and loss are indicated in red and green, respectively (see Table S2). **c.** Genes recurrently amplified or deleted across tumors.

Strikingly, three of five liver tumors from aged p21^⁺/TertCi^ mice (LT981, LT20-3, and LT24) harbored the same activating *Hras* mutation (*Hras^Gln61Lys^*) (Figure 2A) a known hotspot in diethylnitrosamine (DEN)-induced murine liver tumors (Connor *et al*, 2018). LT981 also carried a missense mutation in Rnf43 (*Rnf43^Thr208Pro^*), an E3 ubiquitin ligase. LT20-3 harbored multiple mutations, including a C-terminal deletion in the histone methyltransferase *Kmt2d* (*Kmt2d*^Gln4282del)^, which mediates H3K4 monomethylation at enhancers (Lee *et al*, 2013), a frameshift mutation in *Fga* (*Fga*^Ala245fs^), and a missense mutation in Ep300 (*Ep300*^Ala74Glu^). LT20-1 which arose in the same mouse as LT20-3 harbored the same *Ep300*^Ala74Glu^ mutation but lacked the *Hras^Gln61Lys^* mutation (Figure 2A). LT20-1 also carried a mutation in *Ppp1ca* (*Ppp1ca^Arg191Trp^*), the catalytic PP1 subunit, as well as a frameshift mutation in *Uri1* (*Uri1^Ser481fs^*), a transcriptional repressor implicated in ROS suppression (Xu *et al*, 2021). LT24, which showed extensive genomic rearrangement (Figure 2B) had a truncating mutation in *Ogg1* (*Ogg1^Cys253Ter^*), involved in oxidative DNA damage repair (Boiteux & Radicella, 2000). LT984, classified as a hepatocellular adenoma also carried the *Ep300^Ala74Glu^*mutation, marking as the third tumor with this recurrent mutation. It also harbored a deletion in *Rnf168* (*Rnf168^Glu159Del^*), a DSB-repair E3 ligase and a missense mutation in *Fn1* (*Fn1^Cys335Gly^*), which encodes fibronectin (Figure 2A).

Interestingly, mutations in PP1 subunits were identified in multiple tumors, *Ppp1ca* in LT20-1, *Ppp1r9a* in LT992, and *Ppp1r1a* in LT941 (Figure 2A, Table S1), highlighting a potential role of the PP1 pathway dysregulation in HCC development (see discussion).

### Shared chromosomal alterations between p21^⁺/Tert^ and p21^⁺/TertCi^ tumors mirror human HCC

We analyzed copy number alterations (CNAs) across the different tumors (Figure 2B, Table S2, Fig. EV1). No detectable gains or losses were found in LT992 and LT984 and minimal alterations in LT20-1, and LT20-3 in contrast of all other tumors (Fig. EV1). Recurrent amplifications included a Chr 15 gain in LT662, LT663, LT981, and LT24, a ChrX gain in LT663 and LT941, and a Chr5 gain in LT941 and LT981 (Figure 2B). With respect to chromosomal losses, LT662, LT981, and LT24 showed a loss on Chr 4, often accompanied by a loss on Chr X. A similar pattern of Chr4 and ChrX loss was also detected in the WT liver neoplasm (WT4) (Figure 2B). Interestingly, LT663 showed extensive rearrangement of Chr 14 (Fig. EV1), indicative of chromothripsis, a catastrophic chromosomal shattering and reassembly event previously reported approx. 5% of human HCC cases (Fernandez-Banet *et al*, 2014; Chen *et al*, 2024).

We compiled a curated list of genes recurrently amplified or deleted in human HCC (Zucman-Rossi *et al*, 2015; Schulze *et al*, 2016; Niu *et al*, 2016; Letouzé *et al*, 2017; Yang *et al*, 2023; Chen *et al*, 2024) and found strong parallels in the mouse tumors (Figure 2C). In particular, the conserved amplification on Chr 15 includes several key genes (*Myc, Cthrc1, Rad21, Ndrg1*, and *Ptk2*) linked to liver tumorigenesis, tumor growth, metastasis, or immune evasion (Dhanasekaran *et al*, 2022; Sun *et al*, 2024; Pang *et al*, 2024; Tang *et al*, 2023; Fan *et al*, 2019). Additional amplifications include *Met* in LT663 and *Ccnd1* and *Fgf15* (the murine ortholog of human *FGF19*) in LT24 frequently amplified in human HCC (Schulze *et al*, 2015). Chr 5 gains in LT941 and LT981 includes *Sdhc* and *Parp1*, genes implicated in HCC progression (Bai *et al*, 2022; Paturel *et al*, 2022). Both tumors also showed *Mdm4* amplification, a negative regulator of TP53 found overexpressed in fibrolamellar HCC, particularly in adolescent patients whose tumors often lack TP53 mutations (Karki *et al*, 2019) (Figure 2C). Among genes affected by copy number loss, only LT24 harbors a *Tp53* deletion. *Cdkn2a*, *Cdkn2b*, and *Ard1a* (Chr 4) on their side are deleted in LT662, LT981, and LT24 while LT663, LT981, and LT24 share a deletion of cell cycle regulators *Cdc25* and *Rb1*. Loss of these genes is linked to senescence bypass and liver tumorigenesis (Niu *et al*, 2016; Zucman-Rossi *et al*, 2015; Joseph *et al*, 2019; Reimann *et al*, 2024). LT941 and LT24 also show deletions in alcohol dehydrogenase genes, including *Adh4*, whose loss correlates with poorer HCC prognosis (Zhang *et al*, 2023) (Figure 2C). Unlike human HCC where *TERT* is frequently activated, about half of the tumors exhibited loss of the endogeneous *Tert* locus, likely due to the mouse models used in which *Tert* or *Tert*^Ci^ are expressed under the control of the *p21* promoter, rendering activation of the native locus dispensable.

In conclusion, despite the limited sample size, our findings show that both *p21^+/Tert^*and *p21^+/TertCi^* mouse models-which mimic telomerase activation in pre-senescent cells-develop HCCs with key molecular and histopathological features of human liver cancer.

### Activating RAS mutations in p21^⁺/TertCi^ tumors are associated with elevated mutagenesis

We identified recurrent mutations in *Hras^Gln61Lys^* (LT981, LT24, LT20-3) and *Ep300^Ala74Glu^* (LT984, LT20-1, LT20-3) in p21^+/TertCi^ tumors. Notably, in a (DEN)-induced mutagenesis model, half of all liver tumors harbored *Hras* mutations at position Gln61, with Hras^Gln61Arg^ being the most frequent (Connor *et al*, 2018). We examined the number and pattern of mutations observed across tumors (Figure 3A). Among p21^+/Tert^ tumors, LT662, LT663, and LT941 exhibited high levels of mutations while LT992 was sparsely mutated (Figure 3A). p21^+/TertCi^ Tumors harboring the *Hras^Gln61Lys^* mutation exhibited the higher number of mutations, whereas the two tumors (LT984 and LT20-1) with the Ep300^Ala74Glu^ displayed a lower mutation burden. Mutational signatures analysis in p21^+/Tert^ and p21^+/TertCi^ tumors revealed that C>A transversions (20% of the mutations) occurred at similar overall frequencies in both tumor types while C>T transitions were slightly more frequent (35% et 45% in *p21^+/Tert^* and *p21^+/TertCi^* tumors, respectively) (Figure 3B). C>T transitions occurred preferentially in similar contexts A[C>T]G and C[C>T]G in both type of tumors (Figure 3C). Interestingly, in p21^+/Tert^ tumors C>A transversions predominantly occurred in A[C>A]T and T[C>A]T contexts whereas in p21^+/TertCi^ tumors they were mainly enriched in G[C>A]A and T[C>A]T contexts (Figure 3C). The oncogenic mutations Egfr^Ser303Tyr^ (TCA /TAT) and Ep300^Ala74glu^ (GCA/GAA) arose in such context. At the individual tumor level C>A transversions were enriched in LT941, LT981, LT24, and LT203, all harboring activating RAS mutations (Figure 3D). Oddly LT992 showed no evidence of mutagenesis apart from a few isolated T>G. The few mutations in LT992 are in frame deletions or frameshift variant.

**Figure 3.**
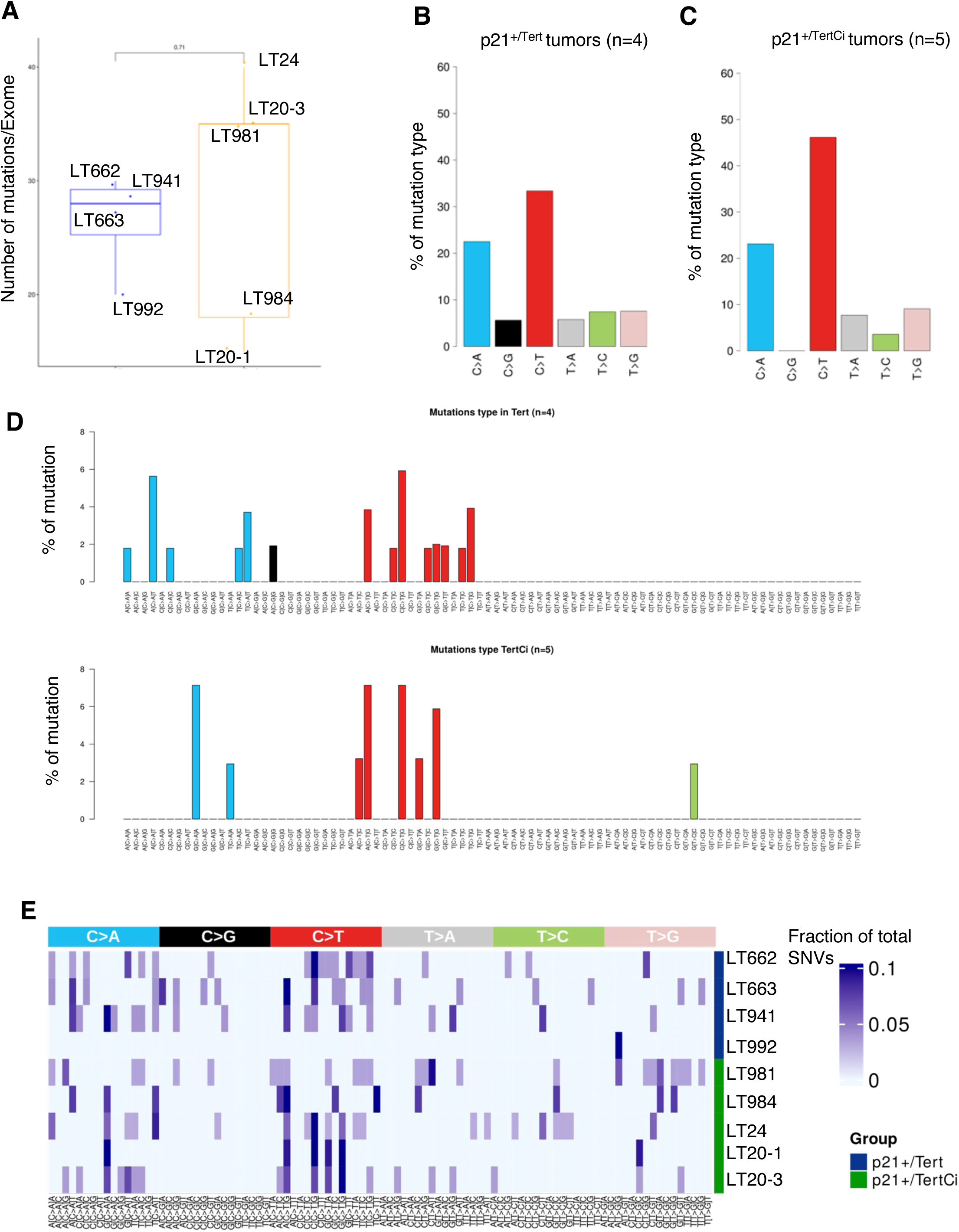
Mutational signature analysis of p21^⁺/Tert^ and p21^⁺/TertCi^ tumors. **a.** Number of somatic mutations per exome in the tumors. **b and c.** Mutation type in p21^⁺/Tert^ and p21^⁺/TertCi^ tumors. **d.** Occurrence of mutations from p21^⁺/Tert^ and p21^⁺/TertCi^ tumor samples classified by substitution type and trinucleotide context. **e.** Heat map showing mutational profiles of individual mouse tumor samples (rows), categorized by substitution type and trinucleotide context (columns).

The mutagenic effect associated with the expression of TERT and TERT^Ci^ and their subsequent amplification though the emergence of oncogenic mutations are addressed in the discussion.

### Oncogenic signalling pathways in p21^+/Tert^ and p21^+/TertCi^ tumors

We performed RNA-seq analysis on nine liver tumors, including four from p21^+/Tert^ mice, five from p21^+/TertCi^ mice, one neoplasm from a wild-type (WT) mouse, and six healthy liver samples collected from tumor-bearing mice. Principal Component Analysis (PCA) revealed a separation between healthy and tumor samples based on their global gene expression profiles (Fig. EV2A), highlighting the distinct transcriptional landscape of the tumors. We identified differentially expressed genes (DEGs) between tumor and healthy liver tissues (Table S3). We found that many of the significantly up- and down-regulated genes were also deregulated in HCCs from the *Axin1Δ* mouse model (Abitbol *et al*, 2018) (Fig. EV2B) and in human HCC, underscoring the biological relevance of our models and the presence of conserved HCC markers across species.

We next analyzed the 200 most differentially expressed genes in either p21^+/Tert^ or p21^+/TertCi^ tumors compared to non-tumoral liver tissues (Figure 4A). Among these, 45 and 25 genes were commonly upregulated or down regulated, respectively, in both tumor types. 155 genes were selectively overexpressed in either p21^+/Tert^ or p21^+/TertCi^ tumors whereas 175 genes were down-regulated. Gene Ontology analysis of Biological Processes (GOBP) associated with the upregulated genes revealed distinct patterns between the two models. In p21^+/Tert^ tumors, enriched pathways were primarily related to sister chromatid segregation and mitosis (Figure 4B). In contrast, p21^+/TertCi^ tumors showed a significant upregulation of genes involved in wound healing, angiogenesis, and the ERK1/ERK2 signaling cascade, aligning with the presence of the *Hras^Gln61Lys^* mutation in three of the five tumors. Consistent with these findings, p21^+/Tert^ tumors overexpress cyclin-dependent kinase 1 (Cdk1) and Cdk6, along with multiple cyclins (Ccna2, Ccnb1, Ccnb2, Ccnd1), in contrast to p21^+/TertCi^ tumors for which such upregulation is in most case not significant (Figure 4C).

**Figure 4.**
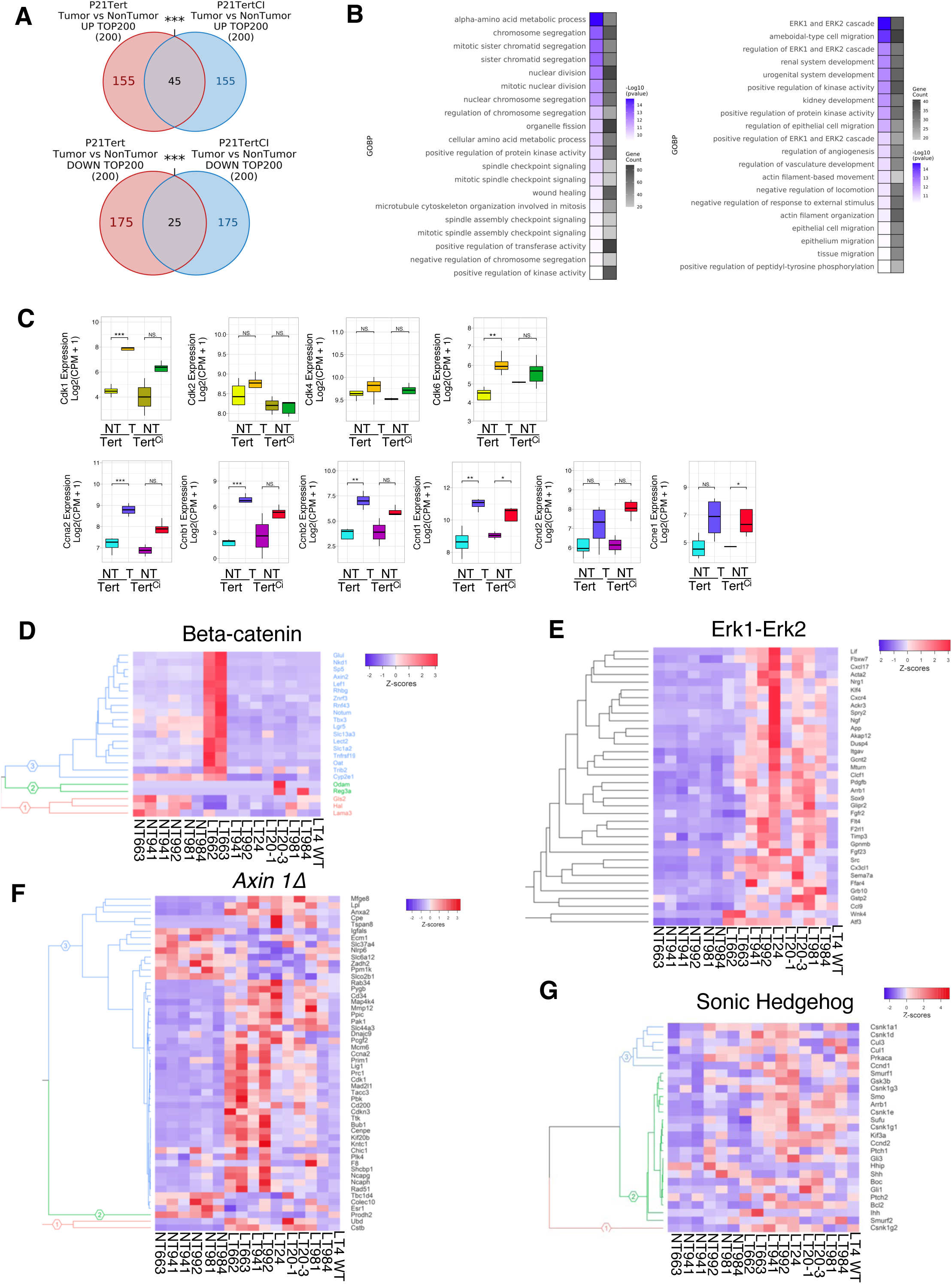
Transcriptomic analysis of p21^+/Tert^ and p21^+/TertCi^ tumors. **a.** Venn diagram showing the number and overlap of significantly (p<0.05) upregulated genes (upper panel) or down-regulated (lower panel) between p21^+/Tert^ and p21^+/TertCi^ tumors. **b.** Gene Ontology Biological Process (GOBP) terms enriched among differentially expressed genes (DEGs) in p21^+/Tert^ (left) and p21^+/TertCi^ (right) tumors. **c.** Expression levels of cyclin dependent kinases and cyclins in p21^+/Tert^ and p21^+/TertCi^ tumors. Statistics *p<0.05, **p<0.01, *p<0.001 vs NT **d-g.** Heatmaps showing the expression profiles of pathway-specific gene signatures, including activated β-catenin, MAPK–ERK, *axin1Δ*, and activated Sonic Hedgehog pathways, in p21^+/Tert^ and p21^+/TertCi^ tumors. Beta-catenin and *axin1Δ* signatures were defined in Abitbol et al (2018).

We next sought to identify transcriptional signatures indicative of signaling pathway activation, specifically Wnt/β-catenin, ERK1/ERK2 (MAPK), Notch, and Sonic Hedgehog (SHH), all of which are frequently implicated in human HCC (Zucman-Rossi *et al*, 2015). As expected from their mutational profiles (*Ctnnb1^Asp32Tyr^* and *Ctnnb1^Thr41Ile^*), LT662 and LT663 displayed strong activation of the Wnt/β-catenin pathway (Figure 4D). These two tumors showed high expression of a ten-gene mutated β-catenin gene signature comprising AXIN2, GLUL, LGR5, NKD1, NOTUM, RHBG, SLC13A3, SP5, TCF7, and TNFRSF19 which has been reported to identify CTNNB1-mutated HCC (Lehrich *et al*, 2024). In addition, LT662 and LT663 showed marked overexpression of the Na⁺-dependent amino acid transporters SLC1A5, which mediates glutamine uptake and has recently been recognized as a novel hallmark of HCC (Tambay *et al*, 2024), and SLC1A2, which transports glutamate.

Tumors harboring the *Hras^Gln61Lys^* mutation, LT24, LT20-3, and LT981 (all from p21^+/TertCi^ mice), along with LT941 (p21^+/Tert^, carrying Egfr^Ser306Tyr^) exhibited a shared transcriptional signature consistent with ERK1/ERK2 (MAPK) pathway activation, in alignment with their mutational backgrounds (Figure 4E). Further, LT662, LT663, and LT992 exhibited transcriptional features of Notch pathway activation, similar to those observed in *AxinΔ* mouse models (Villanueva *et al*, 2012; Abitbol *et al*, 2018) (Figure 4F). A distinct Sonic Hedgehog (SHH) signature was found in LT981, LT984, LT992, and LT24 (Figure 4G), indicating additional heterogeneity in pathway activation across tumors. Overall, the transcriptomic profiles of the tumors are largely consistent with their underlying mutational landscapes. Interestingly, LT992 showed a MAPK activation signature, despite lacking canonical *RAS* pathway mutations. We hypothesize that this activation could result from mutations in phosphatase genes (PtprE and Ppp1r9a), which may lead to increased substrate phosphorylation and thereby activate MAPK signaling (Chen *et al*, 2024).

Among model-specific transcriptional signatures, only p21^+/Tert^ tumors exhibited overexpression of the helicase Pif1 and Ten1, a component of the CST complex (Ctc1/Stn1/Ten1) involved in telomere replication and stability (Wang *et al*, 2012) (Fig. EV3A). Additionally, nearly all tumors, except LT984 and LT20-1, showed strong overexpression of the maternally imprinted long noncoding RNA *H19* and its associated microRNA, miR-675, both of which have been implicated in liver cancer pathogenesis (Wang *et al*, 2023) (Fig. EV3B).

### Tumors exhibit suppressed gluconeogenesis but upregulate distinct Nfrf2 target genes

A recent study (Gu *et al*, 2025) showed that during the progression from metabolic dysfunction–associated steatohepatitis (MASH) to HCC, FBP1 expression declines in premalignant hepatocytes. To explore metabolic alterations in the tumor samples, we analyze the expression of genes involved in glycolysis and gluconeogenesis. Enzymes involved in gluconeogenesis Aldob, Fbp1, Pcx, Pck1, and G6pc (Group 1) were consistently downregulated across all tumors, except for LT984 (Figure 5A). Conversely, glycolytic enzymes (Group 2) were generally overexpressed in all tumors again with LT984 as the exception. In particular, Hk2, Eno1, Tpi1, and Gapdh (Group 3) were strongly upregulated in tumors LT662, LT663, LT201, and LT203 (Figure 5A). These findings indicate that, apart from LT984 (HCA), all tumors suppress gluconeogenesis in favor of enhanced glycolytic activity (Figure 5A). In particular, Fbp1 was downregulated in all tumor samples except LT984 (Figure 5B). Because Fbp1 down regulation was associated to NRF2 activation (Gu *et al*, 2025), we examine the expression of Nrf2 (Nfe2l2) target genes (Malhotra *et al*, 2010; Mitsuishi *et al*, 2012b). We found that most Nrf2 targets involved in redox and energy metabolism regulation (Luchkova *et al*, 2024) were differentially regulated in tumor samples (Figure 5C). Interestingly, p21^⁺/Tert^ and p21^⁺/TertCi^ tumors exhibited distinct Nrf2 target expression patterns based on their mutational profiles (Figure 5C). For instance, LT662 and LT663, which harbor activating mutations in Ctnnb1, preferentially upregulated a subset of Nrf2 targets (Group 1) including Tkt (transketolase) and Taldo1 (transaldolase 1) involved in the pentose phosphatase pathway, core regulator of glutathione synthesis and antioxidant defence (Gclm, Sqstm1, Mafg), as well as Nupr1, a stress responsive gene induced under oxidative and metabolic stress and known to be expressed in human HCC (Huang *et al*, 2021; Cai *et al*, 2025) (Figure 5C). In contrast, tumors LT941, LT992, LT24, LT201, LT203, and LT981 predominantly upregulated genes involved in antioxidant defense (Group 2) including Slc7a11, Gpx4, Nqo1, Txnrd1, Fth1, Hmox1, and Gsr, all being regulators of ferroptosis resistance, a cell death pathway driven by lipid peroxidation (Koppula *et al*, 2021; Su *et al*, 2025) (Figure 5C). When considering all tumors, neither Nrf2 nor Tp53 are significantly overexpressed in tumors (Fig. EV3C). Notably, although Nrf2 mRNA levels were higher in LT992 and LT24 compared to LT203 and LT981, the latter two showed the strongest upregulation of Nrf2 target genes. This suggests that Nrf2 protein stabilization may drive its activity in the tumors rather than its transcriptional regulation.

**Figure 5.**
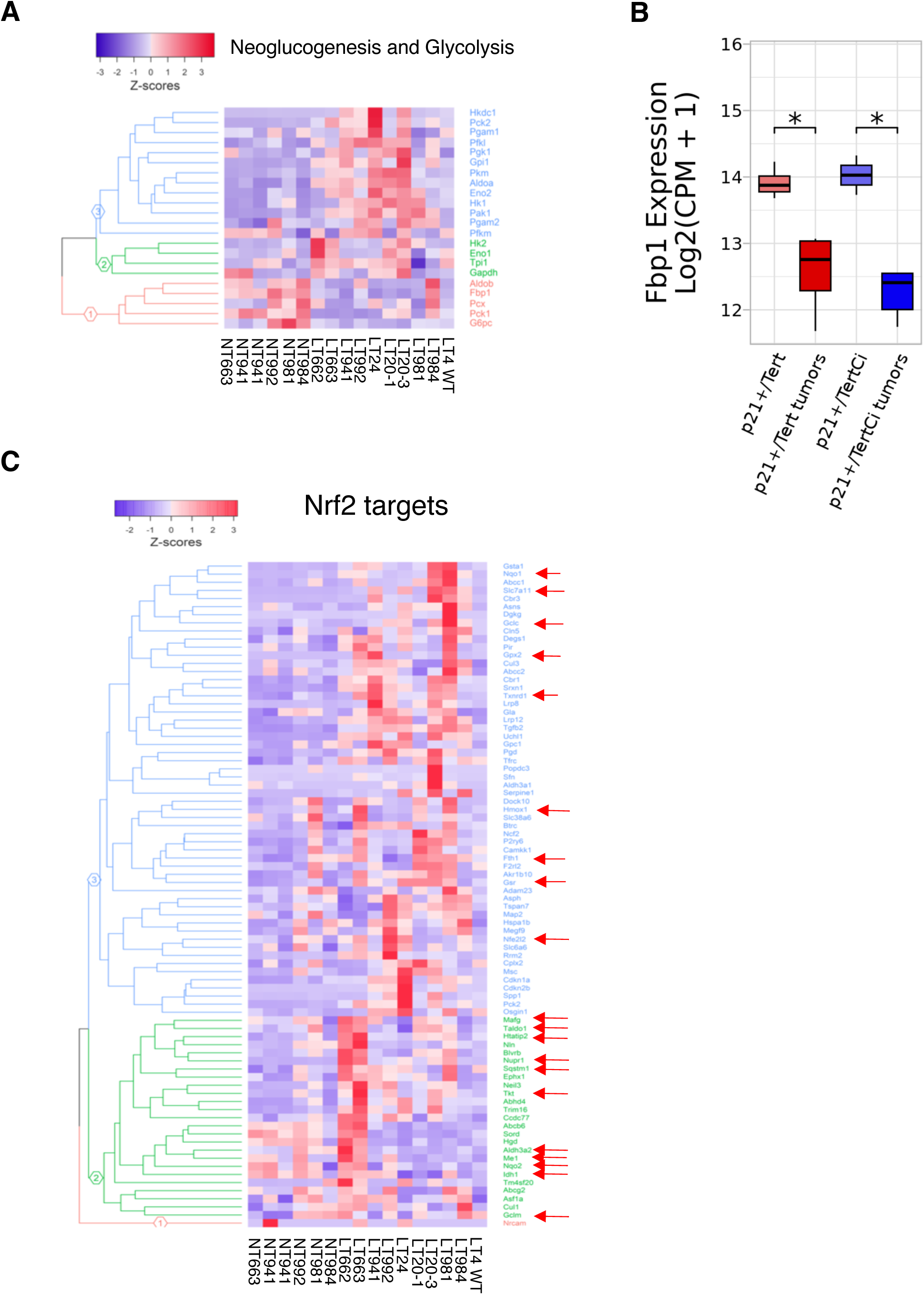
Downregulation of gluconeogenesis and activation of Nrf2 targets in tumors. **a.** Heatmaps showing the expression of genes involved in gluconeogenesis (red group) and glycolysis (Green and Blue groups). **b.** Expression levels of FBP1 in p21^+/Tert^ and p21^+/TertCi^ tumors, Statistics *p<0.05 vs non tumoral p21^+/Tert^ or p21^+/TertCi^ **c.** Heatmaps showing the expression of NRF2 target genes. Genes were clustered in two groups: (green) preferentially expressed in beta-catenin activated tumors, (blue) preferentially expressed in MAPK–ERK activated tumors. Arrows indicate genes enhancing antioxidant capacity and glutathione metabolism thereby protecting cells from ferroptosis.

### Spatial analysis reveals altered HNF4alpha protein levels and liver tissue architecture

We employed the Hyperion™ Imaging System, which enables high-dimensional, subcellular-resolution analysis of multiple cellular targets simultaneously in formalin-fixed paraffin-embedded (FFPE) tissues (Elaldi *et al*, 2021). Using a validated panel of 10 metal-conjugated antibodies (Table S4), we analyzed the architecture of the tissues. To characterize the liver parenchyma, we first stained healthy livers and the tumors from p21^+/Tert^ and p21^+/TertCi^ mice using antibodies directed against beta-catenin (Ctnnb1), Hepatocyte nuclear factor 4α (HNF4alpha), and alpha Smooth Muscle Actin (α-Sma) (Figure 6A, B). Scale bars (200 µm) corresponding to the uncropped images are provided in Fig. EV4. In healthy p21^+/Tert^ (L1069) and p21^+/TertCi^ (L982) livers (Figure 6A), a majority of cells were stained by HNF4alpha antibodies likely marking non-transformed hepatocytes. Beta-catenin displayed a typical wild-type distribution with predominant localization at hepatocyte plasma membrane consistent with its role in cell-cell adhesion and α-SMA was primarily restricted to the perivascular regions where smooth muscle cells reside (Figure 6A). It was challenging to distinguish whether 4T1 or LT662 and LT663 tumors, which share an activating mutation in Ctnnb1, exhibited higher levels of cytoplasmic or nuclear beta-catenin indicative of beta-catenin activation (Figure 6A). Notably, all tumors, showed a reduced number of HNF4alpha-positive cells as well as a decrease in staining intensity (see the following paragraph for HNF4alpha quantification across tumors). Moreover, unlike the perivascular localization observed in the healthy livers, we observed dispersed α-Sma positive cells throughout the tumor likely reflecting the presence of cancer associated fibroblasts (Sun *et al*, 2023; Xu *et al*, 2022). In tumors LT941, LT992, and LT201, α-Sma positive areas accounted for more than 20% of the area occupied by DNA, indicating an important stromal activation.

**Figure 6.**
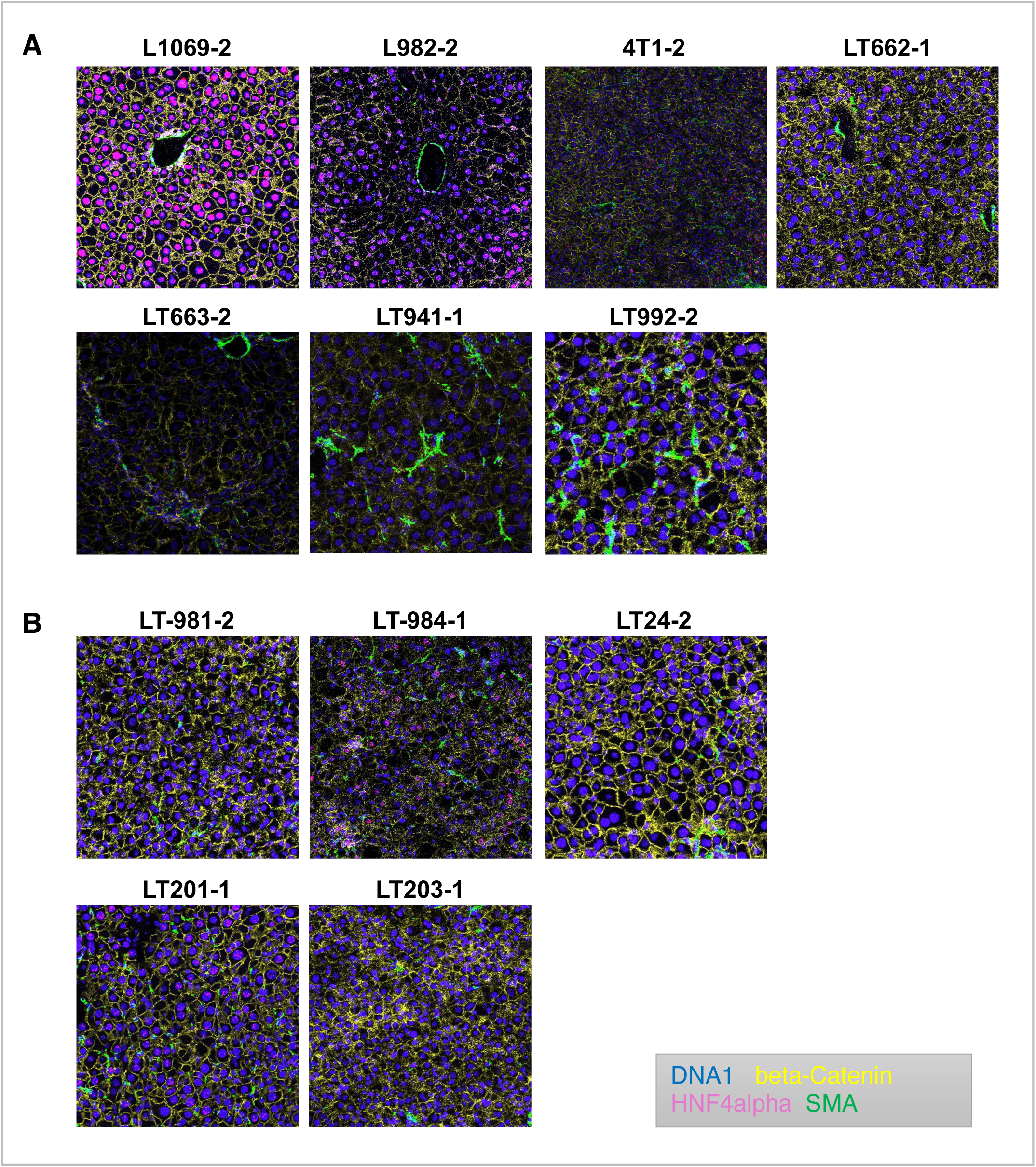
Spatial architecture of liver tumors. Non tumoral and tumor FFPE tissue sections were analyzed using the Hyperion™ Imaging System with antibodies against CTNNB1 (Yellow), HNF4alpha (Magenta), and α-SMA (Green). L1069 and L982 represent to non-tumoral liver tissue from p21^+/Tert^ and p21^+/-^ mice, respectively. Tumors numbered in the figure are described in Table 1. The number following each tumor name indicates the region of interest (ROI). Iridium intercalator was used for nuclear staining. *High resolution, uncropped images are included in the supplemental files*.

We next quantified HNF4alpha and also p21 positive cells across all the samples (Figure 7A, B). In healthy p21^+/Tert^ and p21^+/TertCi^ livers 40 to 50% of liver cells expressed HNF4alpha. This proportion dropped sharply to 10% or less in the tumors harboring the Ctnnb1^Asp32Tyr^, Ctnnb1^Thr41Ile^, Egfr^Ser303Tyr^, and Hras^Gln61Lys^ mutations (Figure 7A, B). In contrast, tumors lacking activation of the Wnt/β-catenin or hRAS signaling pathways retained approximately 20% of HNF4alpha-positive cells (Figure 7A, B). Reflecting hepatocyte dedifferentiation, tumors exhibited strong downregulation of major urinary protein genes, normally produced by hepatocytes in mice (Fig. EV5A) (Krauter *et al*, 1982).

**Figure 7.**
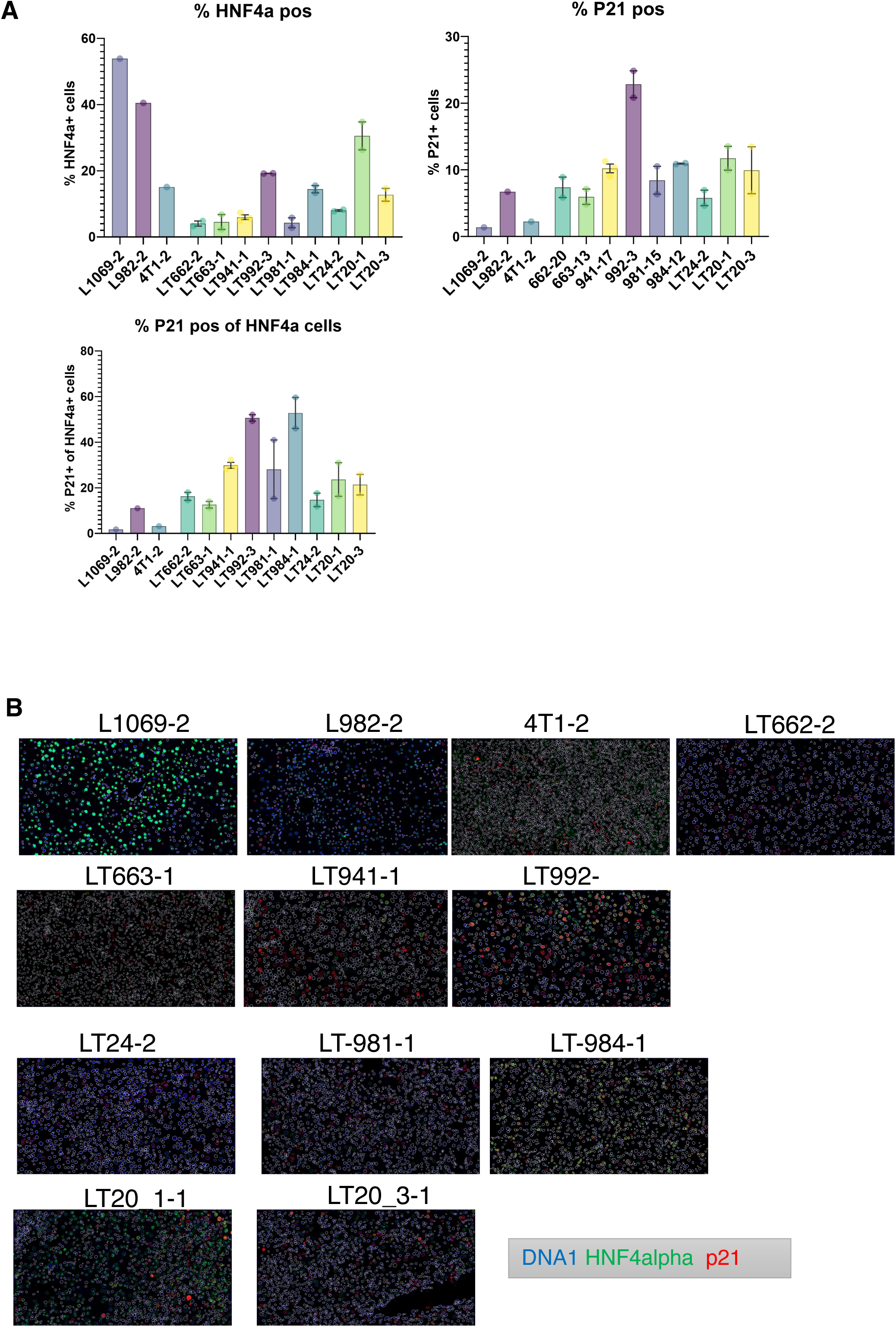
Quantification of HNF4alpha and p21 levels across tumor samples. **a.** ROIs from liver tumor sections were imaged using the Hyperion™ with antibodies against HNF4alpha and p21. Signal quantification was performed as described in the Materials and Methods. The percentage of HNF4alpha, p21, and double-positive cells are shown. The 2 dots for each tumor represent two different ROI. **b.** Representative images of non-tumoral and tumoral sections. HNF4alpha (Green), p21 (Red), and nuclei (Blue). *Mosaic uncropped images are included in the supplemental files*.

These findings prompted us to examine the transcript levels of HNF4alpha. RNA-seq analysis did not reveal significant downregulation of Hnf4alpha mRNA levels across tumor samples (Table S3) suggesting that HNF4alpha may be subject to post-transcriptional regulation in tumors with an activation of the Wnt/β-catenin or hRAS signaling.

On the same FFPE sections, we quantified the proportion of p21-positive cells. In L1069 (p21^+/Tert^) and 4T1 (p21^+/+^) livers, fewer than 2% of cells were p21-positive compared to approximately 5 % in L982 healthy liver (p21^+/TertCi^) (Figure 7A, B). This proportion increased to above 10% in LT941, LT992, LT984, and LT201 (Figure 7A, B) and was about 5% for the other tumors consistent with the transcriptional overexpression of p21 in all tumors (Table S3). One tumor (HCA), LT992, exhibited a markedly higher proportion of p21-positive cells (>20%) (Table S3). Interestingly, in LT992 and LT984 both classified as hepatocellular adenoma (Figure 1, Table 1), approximately 50 % of HNF4alpha-positive cells were also positive for p21. Notably, both tumors display high levels of TERT mRNA, suggesting that TERT may drive hepatocyte proliferation in these two tumors which lack mutations in the Wnt/β-catenin and hRAS pathways.

### Tumor infiltration by immune cells is altered in β-catenin–activated HCC

We further assessed immune cell infiltration within the tumors. To characterize the infiltration of lymphocyte classes, we used metal-conjugated antibodies against CD3 (T lymphocytes), CD4 (auxiliary T lymphocytes), CD8 (cytotoxic T lymphocytes), and B220 (B lymphocytes). We found that the two beta-catenin activated tumors (LT662 and LT663) were poorly infiltrated by T-lymphocytes (2 to 4 % of CD3-postive cells) with 1-2% of auxiliary and cytotoxic T lymphocytes (Figure 8A, B). These results are consistent with previous studies showing that activation of the Wnt/β-catenin pathway in HCC is associated with poor T-lymphocyte infiltration, facilitating immune evasion and resistance to anti-PD1 therapy (Dantzer *et al*, 2024). In contrast, LT992, LT201, and LT941 tumor showed higher level of T lymphocyte infiltration (> 10 %) (Figure 8A, B). While LT201 was characterized by a higher proportion of auxiliary (CD4⁺) T cells (8-10 %) compared to cytotoxic (CD8⁺) T cells (<4%), the highly aggressive LT941(Egfr^Ser303Tyr^) displayed a predominance of cytotoxic T cells (approx.10%) over auxiliary T cells (Figure 8A,B). LT981, LT203, and LT24 tumors, all carrying the Hras^Gln161Lys^ mutation, displayed comparable intermediate levels of lymphocyte infiltration (approx.5%). In summary, the two β-catenin–activated tumors exhibited the lowest levels of T-cell infiltration, whereas the Egfr-activated tumor along with tumors LT992 and LT201 (which both harbor mutations in subunits of PPP1 exhibited the highest degree of T lymphocyte infiltration (approx. 5%). Regarding B lymphocyte infiltration, tumors exhibited very low levels of infiltrating B lymphocytes (1 to 3%) (Figure 8A, B). Only LT981 exhibited a proportion of infiltrating B lymphocytes exceeding 5%.

**Figure 8.**
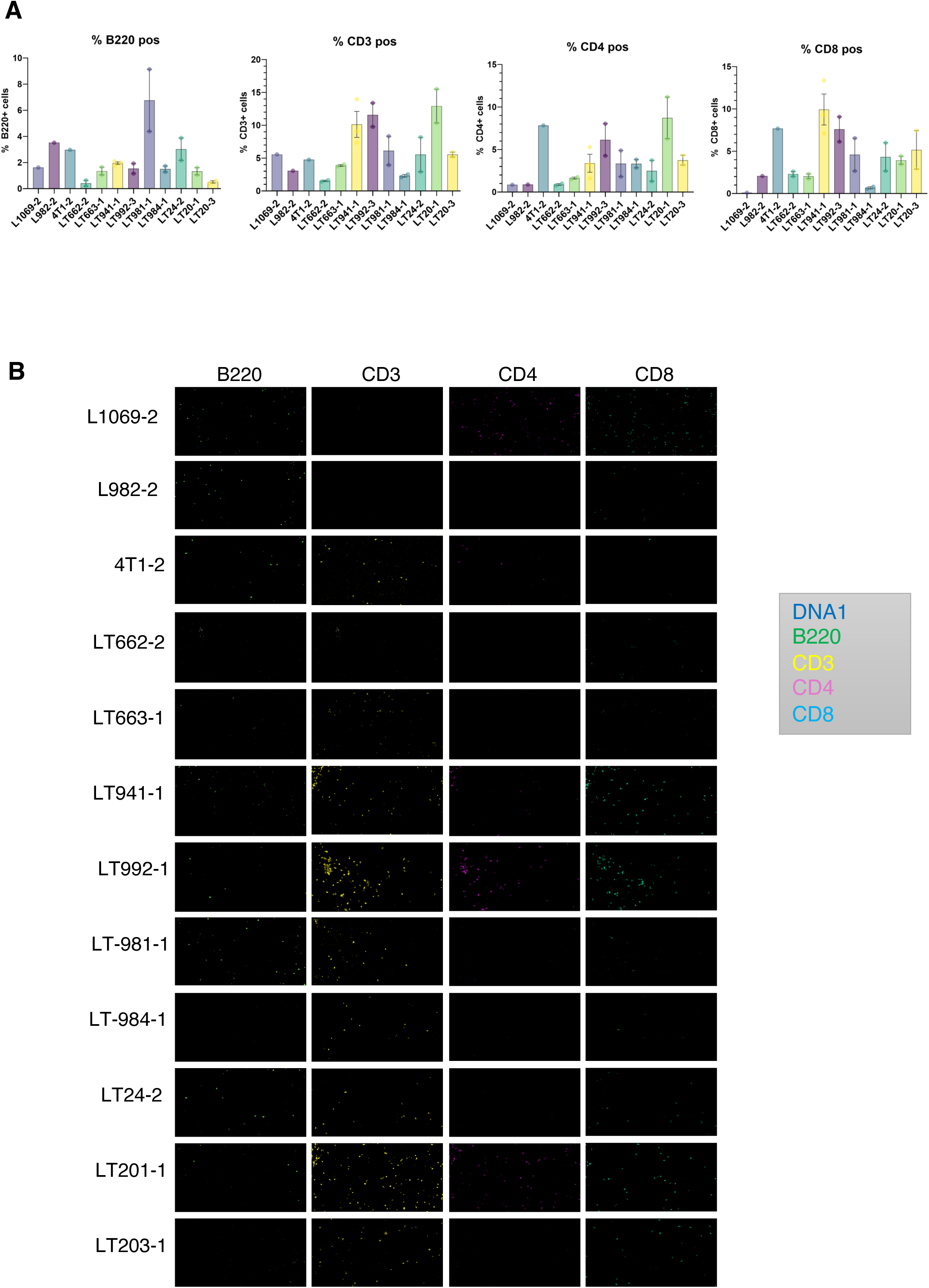
Tumor infiltration by lymphocyte subsets. **a.** ROIs were imaged using antibodies against B220, CD3, CD4, CD4 marking B-lymphocytes, T-lymphocytes, auxiliary (helper) T-cells, and cytotoxic T-cells. Signal quantification was performed as described in the Materials and Methods and immune cell infiltration is shown for each tumor. **b.** Representative images of control a: B220 (Green), CD3 (Yellow), CD4 (Magenta), CD8 (Cyan) and nuclei (Blue). *Mosaic uncropped images are included in the supplemental files*.

We further quantified the proportion of macrophages (F4/80) and dendritic cells (Cd11c) in the liver samples (Figure 9A, C). In healthy liver samples (L1069 and L982), F4/80 positive-cells accounted for approximatively 15% of total liver cells likely, consistent with their identification as kupffer cells (Figure 9A, C). This proportion increase to about 25% in the 4T1 neoplastic tissue carrying the *Ctnnb1^Se45Phe^* but not the activated beta-catenin signature. The percentage of macrophages in tumors ranged from 5% in minimally infiltrated tumors (LT662 *Ctnnb1^Asp32Tyr^*, LT663 *Ctnnb1^Thr41Ile^*, LT984 *Ep300^Ala74Glu^*) to 20-25% in highly infiltrated tumors (LT941 *Egfr^Ser303Tyr^*, LT981 *Hras*^Gln61Lys^, LT203 *Hras*^Gln61Lys^), suggesting that the degree of macrophage infiltration is associated with the oncogenic driver and reflects heterogeneity within the tumor microenvironment (TME). Among these macrophages, in both healthy and tumor tissue of aged p21^+/Tert^ and p21^+/TertCi^ mice, only 2 to 5% were positive for p21indicating that the vast majority of macrophages do not express p21 (Figure 9B, C). Dendritic cells (Cd11c) infiltration was low in healthy livers (1-2%) and in LT662 and LT663 tumors, which show generally weak immune cell infiltration as well as in LT984 (HCA) (Figure 9A, C). LT992, LT24, and LT201 showed higher levels of dendritic cell infiltration with marked punctuate staining (Figure 9C). Interestingly, tumors such as LT941 and LT203 tumors, being markedly infiltrated by macrophages (>20%), exhibited low levels of dendritic cells (3 to 7 %) suggesting competitive dynamics between macrophages and dendritic cells in the TME.

**Figure 9.**
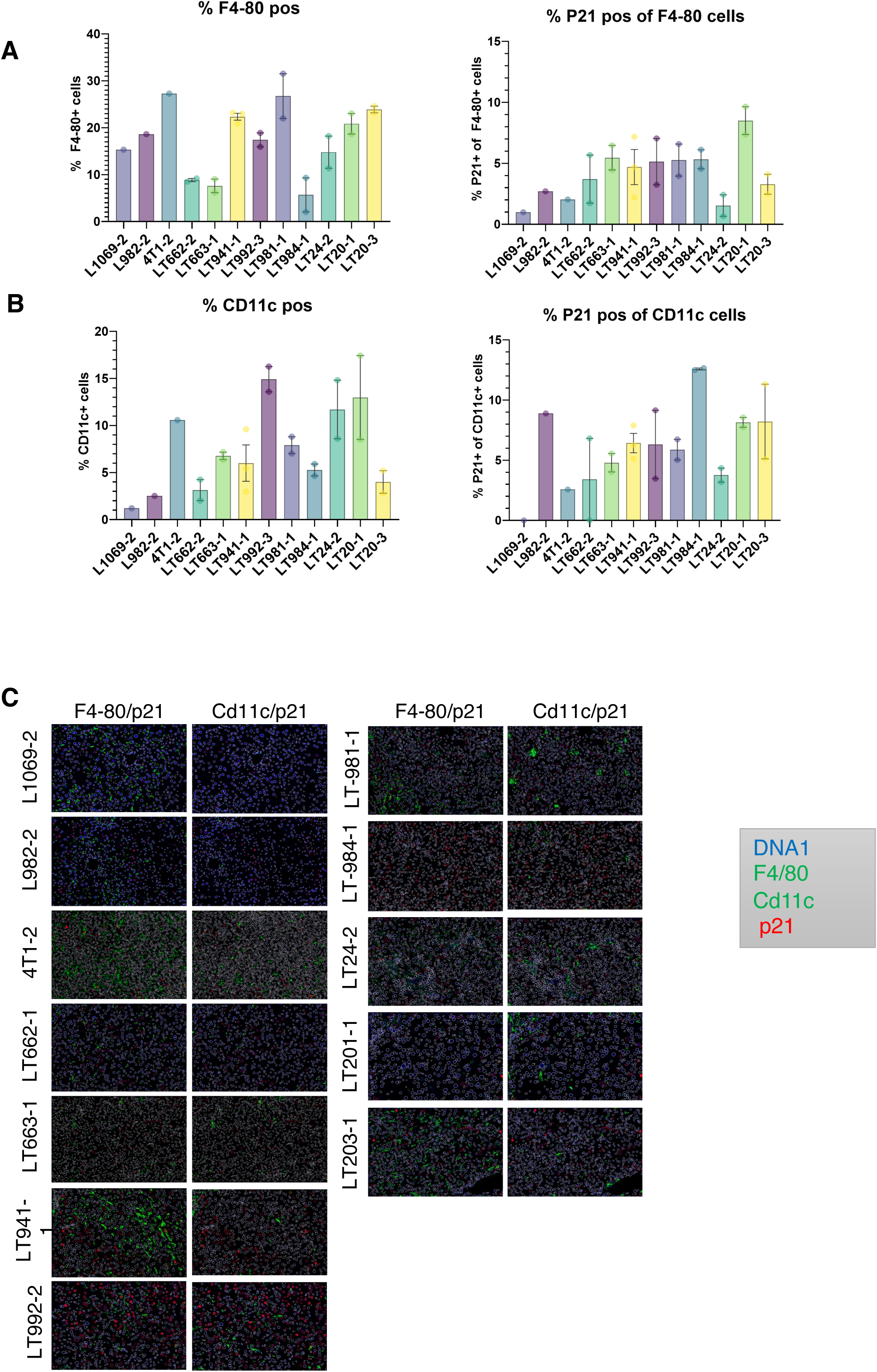
Tumor infiltration by macrophages and dendritic cells. **a.** ROIs were imaged using antibodies against F4/80 and CD11c to label macrophages and dendritic cells, respectively. Signal quantification was performed as described above, and the percentage of positive cells is shown for each tumor. **b.** Representative images of tumor sections. F4/80 is shown in green, CD11c in magenta, and nuclei in blue. *Mosaic uncropped images are included in the supplemental files*.

## Discussion

The p21^⁺/Tert^ and p21^⁺/TertCi^ mouse models develop hepatocellular liver tumors, including hepatocellular carcinomas (HCC) and hepatocellular adenomas (HCA). In our murine models, HCCs resemble human HCCs in many histomorphological aspects, including trabecular and solid growth patterns, multifocal necrosis, hemorrhage, and well- to moderately differentiated tumor cells with mild to moderate atypia. These features support the translational relevance of these models for studying human HCC. However, a notable difference lies in the condition of the adjacent non-tumoral liver: while human HCCs typically arise in a cirrhotic liver background (in 80–90% of cases), the surrounding parenchyma in these murine models exhibits only minimal to mild fibrosis and non-significant liver damage, indicating a divergence in the underlying liver pathology and disease context. It is important to note that by 9 months of age, male p21^+/Tert^ and p21^+/TertCi^ mice developed obesity accompanied by hepatic steatosis, which was associated with altered feeding behavior (L. Braud and V. Géli, unpublished results), suggesting a link between early metabolic dysfunction and later tumor development.

The finding that conditional expression TERT^Ci^ promotes liver tumorigenesis in aged mice raises questions about the mechanism through which TERT^Ci^ drives tumor development. Since models p21^+/Tert^ and p21^+/TertCi^ retain an intact endogenous *Tert* locus, the possibility of endogenous Tert reactivation was considered. However, no mutations were found in the *Tert* promoter region, and CNA analysis suggest a loss rather than amplification of *Tert* locus (Figure 2C). We rather favor the possibility is that TERT^Ci^ modulates a signaling pathway that promotes liver tumor development. This analysis is complex as most tumors are also associated with oncogenic mutations. Comparison of transcriptional profiles between healthy control and p21^+/Tert^ or p21^+/TertCi^ livers from 18-month-old mice reveals that the proto-oncogenes Fos and Jun are overexpressed mainly in p21^+/TertCi^ healthy livers from aged mice, suggesting that p21^+/TertCi^ livers may be in a pro-tumorigenic state that could promote the occurrence of HCC (Bakiri *et al*, 2017, 2024). Another possibility is that TERT^Ci^ exerts a protective capping function at chromosome ends that does not rely on telomere elongation. Supporting this hypothesis, Tpp1 which recruits TERT a chromosome ends (Zhong *et al*, 2012) is expressed across all tumors. Finally, it is also possible that a mitochondrial form of TERT^Ci^ (and TERT) contributes to tumor development by safeguarding mitochondrial DNA and function (Haendeler Judith *et al*, 2009; Saretzki, 2014) in tumors that are exposed to high oxidative stress. This hypothesis is consistent with the observed overexpression of Src in all tumors, as Src kinase activity has been shown to promote the nuclear export of TERT (Haendeler *et al*, 2003). At another level, Pif1 and Ten1 involved in telomere replication and processing (Snow *et al*, 2007; Feng *et al*, 2018; Sanz-Moreno *et al*, 2025) were overexpressed only in p21^+/Tert^ tumors. PIF1 was identified in a study as one of nine prognostic mRNAs in human hepatocellular carcinoma (HCC) (Ni *et al*, 2020) suggesting that overexpression of PIF1 is associated with TERT reactivation.

We observed that p21^⁺/Tert^ tumors were associated with distinct oncogenic mutations. Specifically, the Ctnnb1^Asp32Tyr^ and Ctnnb1^Thr41Ile^ variants are both well-characterized exon 3 gain-of-function mutations known to activate Wnt/β-catenin signaling (Rebouissou *et al*, 2016; Dantzer *et al*, 2024). Consistent with this, LT662 and LT663 displayed a strong transcriptional signature of β-catenin activation (Figure 4D). In addition to β-catenin mutations, LT662 and LT663 carried mutations in Kdm4d (Jmjd2d) and Kdm3c, respectively, two distinct JmjC H3K9 demethylases. Notably, KDM4D has been reported to physically interact with β-catenin and to demethylate H3K9me3 at promoters of β-catenin target genes in colorectal cancer cell lines (Peng et al., 2019). This raises the intriguing question of whether the Kdm4d^Gln73His^ mutation observed in LT662 might impact β-catenin–driven transcription in this context. Notably, mutations *Kdm4d* and *Kdm3c* were exclusive to LT662 and LT663, suggesting potential roles in Wnt pathway modulation. Surprisingly, LT992 exhibited no genomic rearrangements but did harbor two frameshift mutations in the phosphatases *Ppp1r9a* and *PtprE*. Similar mutations were observed in multiple tumors in this study (Figure 2A, Table S1). Of note, recent work has reported mutations in UTR regions of PP1 subunits in human HCC (Chen *et al*, 2024). At the transcriptional level, LT992 displays a distinct signature shared with LT662 and LT663 tumors but distinct to the beta-catenin activated signature that, in both humans and mice, is associated with *AXIN1* mutations (Abitbol *et al*, 2018). PP1 has been shown in human cells to bind AXIN and oppose its phosphorylation (Luo *et al*, 2007). We observed suppressed Wnt signaling in LT992 (Fig. 4D), consistent with the expected effect of PP1 inactivation, which likely increases AXIN1 phosphorylation, enhances GSK3-mediated β-catenin degradation, and thereby dampens Wnt pathway activity (Luo *et al*, 2007). This raises the question of what mechanism drives the shared transcriptional signature observed in both Axin1-mutated HCCs and tumors LT662 (Ctnnb1^Asp32Tyr^), LT663 (Ctnnb1^Thr41Ile^), and LT992 (Ppp1r9a*^Leu1273Thrfs^*) (Fig. 4F). LT992 also carried a truncating mutation in *Sucla2* (Gln87del), previously linked to succinyl-CoA accumulation and protein hypersuccinylation. Elevated protein succinylation has been observed in HCC tissues compared to adjacent normal liver and may be associated with poor prognosis (Bai *et al*, 2022), although its direct role in hepatocarcinogenesis remains uncertain. LT941 also harbored a splice-site variant in another PP1 regulatory subunit, *Ppp1r1a*, along with a mutation in *Ptpn5* (Arg246Gly), which encodes protein phosphatase 5 (PP5). PP5 inhibition has been shown to suppress HCC growth by activating AMPK signaling (Chen *et al*, 2017). Interestingly, LT941 also exhibited the AXIN1-associated transcriptional signature, further reinforcing the connection between phosphatase subunit mutations and AXIN1-related pathway dysregulation. These findings underscore the significant potential role of phosphatases in liver cancer development.

Surprisingly, three of five tumors from aged p21^⁺/TertCi^ mice (LT981, LT20-3, LT24) shared the activating *Hras*^Gln61Lys^ mutation. Gln61 is a mutation hotspot (*Gln61Lys*, *Gln61Arg*, *Gln61Leu*) found at high frequency in DEN-induced liver tumors, but absent in spontaneous cases (Connor *et al*, 2018). LT981, LT20-3, LT24 displayed the highest level of mutagenesis. The finding that tumors harboring activating RAS mutations exhibit increased mutagenesis was notably evident in the comparison between LT20-1 and LT20-3 which originate from the same liver and are genetically similar except for the presence of *Hras^Gln61Lys^* in LT20-3. Of note, the Hras^Gln61Lys^ (CAA/AAA) arose from a C>A transversion. In the DEN-induced mouse model, T>A and T>C predominate reflecting the mutagenic properties of DEN (Connor *et al*, 2018). In contrast, tumors from the p21^⁺/Tert^ and p21^⁺/TertCi^ mice show a higher frequency of C>A and C>T mutations. The predominance of C>T transitions in ACG and CCG contexts (Fig. 3e) suggests that the enrichment of C>T mutations in both tumor types may reflect proliferation-associated cytosine deamination likely at CpG dinucleotides (Alexandrov *et al*, 2020; Tate *et al*, 2019) although LT941 (Egfr^Ser303Tyr^) despite being is the most highly proliferative tumor did not exhibit higher overall mutational burden compared with *Hras^Gln61Lys^*mutated tumors (Fig. 3a). On their side, C>A transversions are often linked to oxidative DNA damage. Notably, these transversion occurred predominantly in distinct trinucleotide context (ACT and GCA in p21^⁺/Tert^ and p21^⁺/TertCi^ mice, respectively), both contexts being consistent with oxidative stress, but the difference may be linked to the associated oncogenic mutation reflecting mechanistic difference in tumor initiation and evolution in these models. Interestingly, LT24 which exhibits the highest mutational burden and mutations across multiple nucleotides carries a truncating in Ogg1^Cys253Ter^ mutation. The results raise the question of whether *TERT^Ci^* expression contributes to the emergence of the Hras*^Gln61Lys^* (CAA/AAA) mutation through a direct or indirect specific mutagenic effect, or whether it instead reflects enhanced cell survival and proliferation. In our previous work, we found that TERT, but not TERT^Ci^, reduces oxidative damage (Lipskaia *et al*, 2024; Braud *et al*, 2025). The high mutation burden in *Hras*-driven tumors might be a consequence of the oncogene’s own mutagenic activity. Similarly, LT984, LT20-1, and LT20-3 all harbor the same *Ep300* mutation (Ala74Glu; GCA→GAA), which also arises from a C>A transversion. *EP300* mutations are present in only 2% of hepatocellular carcinoma (HCC) cases in the TCGA Pan-Cancer Atlas cohort (n = 366, whole-exome sequenced). The functional significance of the Ala74Glu variant remains unknown. As mentioned in the results LT20-1 harbors a mutation in Ppp1ca while LT20-3 harbors a mutation in Kmt2d. *KMT2D* is one of the genes frequently disrupted by hepatitis B virus (HBV) integration in human HCC (Sung *et al*, 2012), and mutations are commonly seen in DEN-induced HCC (Connor *et al*, 2018). p21^+/TertCi^ harbors several additional mutations. Notably, LT981 and LT984 carry mutations in two distinct E3 ubiquitin ligases: *Rnf43* and *Rnf18*, respectively. *RNF43* mutations are frequently observed in colorectal and gastric cancers (Giannakis *et al*, 2014; Wang *et al*, 2014), and have been associated with poor prognosis in liver cancer (Belenguer *et al*, 2022). *RNF168* targets the histone demethylase KDM4A which specifically demethylates H3K9me3 and H3K36me3, for proteasomal degradation (Mallette *et al*, 2012). Loss of *RNF18* may therefore lead to KDM4A stabilization, potentially altering histone methylation dynamics and epigenetic regulation. In addition, LT984 harbored a mutation in *Fn1* (*Fn1^Cys335Gly^*), encoding fibronectin. *FN1* is often affected by HBV integration in non-tumor liver tissues adjacent to HCC (Sung *et al*, 2012) but its direct role in tumorigenesis remains unclear.

Although p21^+/Tert^ and p21^+/TertCi^ have distinct mutations in oncogenic drivers, they exhibit similar patterns of copy number alterations. For example, LT662 and LT663 share amplifications in Chr15 with LT981 and LT24, which are associated with tumor development. However, these four tumors display distinctly different transcriptional profiles, suggesting that oncogenic mutations play a major role in shaping their transcriptional landscapes. Notably, LT992, LT20-1, and LT20-3 exhibit no or little CNA, reinforcing the idea that driver mutations primarily shape transcriptional patterns. Interestingly, half of the HCCs exhibit deletions in various genes involved in cell cycle regulation, including *Cdkn2a*, *Tp53*, and *Rb1*. Loss of these genes are likely to facilitate the bypass of cellular senescence.

Transcriptomic analysis reveals that all tumors except LT984, identified as a hepatocellular adenoma, exhibit downregulation of *Fbp1*. This reduction has been associated with activation AKT and NRF2 pathways, along with decreased expression of p53 and p21, collectively facilitating the bypass of cellular senescence and accumulation of somatic mutations in transformed hepatocytes (Gu *et al*, 2023, 2025). Consistent with these findings, we observed overexpression of *Nrf2* target genes in both p21^+/Tert^ and p21^+/TertCi^ tumors. However, this upregulation did not correlate with *Nrf2* mRNA levels, suggesting that stabilization of the NRF2 protein, rather than increased transcription, may underlie the activation of its targets (Zhu *et al*, 2025). Importantly, LT662 and LT663, which harbor activating mutations in *Ctnnb1*, display a distinct pattern of *Nrf2* target gene upregulation. These upregulated genes are involved in NADPH generation, red-ox regulation, and oncogene-induced stress such as Nupr1. The NRF2 signature in HRAS-mutated tumors exhibits greater heterogeneity, indicating more complex activation of the pathway. Overexpressed genes include NRF2 targets involved in glutathione biosynthesis, reactive oxygen species (ROS) detoxification, and ferroptosis resistance including Slc7a11, features consistent with the increased oxidative stress specific to RAS-induced tumors. It has recently been shown that SLC7A11 is significantly overexpressed in hepatocellular carcinoma (HCC) and that its high expression correlates with lower overall survival, thus highlighting its prognostic value (Zhang *et al*, 2025). Pan-cancer analysis revealed that elevated SLC7A11 associates with higher tumor mutation burden and immunosuppressive microenvironment features (Zhang *et al*, 2025).

Spatial characterization of tumors shows that HNF4alpha levels are greatly reduced in HCC with activation of the beta-catenin and hRAS pathways. HNF4alpha is a central regulator of liver function and acts as a tumor suppressor in HCC (Kotulkar *et al*, 2024). It plays a pivotal role in hepatic glucose metabolism (Shokouhian *et al*, 2023) and contributes to the maintenance of hepatocyte quiescence, in part by repressing *cyclin D1* (Ccnd1) expression (Wu *et al*, 2020). Consistently, Ccnd1 was markedly overexpressed in the tumors. In line with findings that identify HNF4alpha as a key regulator of the transition from slow-growing tumors to aggressive HCC (Lazarevich *et al*, 2010), 4T1, LT992 (HCA), and LT984 (HCA) cell lines exhibited intermediate levels of *Hnf4alpha* expression. Surprisingly, we observed that HNF4 alpha levels were downregulated at the post-transcriptional level. In human colon cancer, phosphorylation of HNF4alpha by Src tyrosine kinase has been associated with the loss of specific isoforms of HNF4alpha (Chellappa *et al*, 2012). Degradation of HNF4alpha was further confirmed in liver (Huck *et al*, 2019). As noted above, Src was upregulated in all tumors, albeit to a lesser extent in LT662 raising the possibility that kinases, such as Src may promote HNF4alpha degradation in these tumors.

We next analyzed immune cell infiltration in two distinct regions of the tumor. Despite the small size of these regions (1-2 mm²), the analysis revealed comparable patterns of immune infiltration across both areas of the tumors. Constitutive activation of β-catenin has been shown to suppress immune cell recruitment (Lehrich & Monga, 2025). Approximately 70% of WNT/β-catenin–active tumors are classified as immune-excluded, characterized by low T-cell infiltration (Montironi *et al*, 2023). It has been shown that the immune evasion associated with beta-catenin activation observed in both murine and human HCC results from impaired dendritic cell recruitment (Ruiz de Galarreta *et al*, 2019). Consistent with these findings, LT662 and LT663, which exhibit high expression of AXIN2 and GLUL, exhibited low infiltration by T lymphocytes as well as by macrophages and dendritic cells. Chemokines known to attract T-lymphocytes (Ccl2, Ccl5, Ccl17, Ccl22, Cxcl9, Cxcl10, and Cxcl11) showed variable levels of expression across all tumors and their levels could not explain the reduced T lymphocyte infiltration in LT662 and LT663. Only Cxcl12 was consistently down-regulated in these two tumors as well as in several other HCC. The observed immune exclusion may therefore reflect, at least in part, tumor-specific adaptations in metabolic reprogramming and oxidative defense.

Interestingly, Glul-overexpressing tumors LT662 and LT663 specifically exhibited a ∼20-fold and a ∼10-fold upregulation of *Slc1a5*, whose gene product transports glutamine into cancer cells (Bhutia *et al*, 2015) and of the glutamate transporter *Slc1a2,* respectively. We predict that LT662 and LT663 establish a glutamine- and glutamate-depleted tumor microenvironment by combining high-affinity uptake transporters (SLC1A2, SLC1A5) with intrinsic glutamine synthesis (GLUL), thereby achieving metabolic autonomy while starving immune and stromal cells of critical nitrogen sources. Recently, glutamine was shown to function as a metabolite regulating the cross-talk between tumors and conventional type 1 dendritic cells (cDC1s), thereby licensing cDC1s to activate cytotoxic T cells (Guo *et al*, 2023). It may therefore that tumor evasion in LT662 and LT663 results from reduced glutamine in the TME (Fig. EV5B). The elevated intracellular glutamine concentration may also promote increased glutathione production and strengthen antioxidant defences. Whether this principle extends to human β-catenin–activated tumors remains to be determined. Recent analyses in human HCC revealed that SLC1A5 expression was associated with poor prognosis and an immunosuppressive microenvironment providing a marker to predict immunotherapy response (Zhang *et al*, 2024a, 2024b). hRAS-activated tumors on their side exhibit preferential upregulation of SLC7A11 that imports cystine in exchange for glutamate to support antioxidant defense and protection against ferroptosis (Li *et al*, 2024).

## Materials and Methods

### Animals

Wild-type (C57BL/6J N), p21^+/Tert^, and p21^+/TertCi^ mice were housed under controlled environmental conditions: temperature (21 ± 1 °C), humidity (60 ± 10%), and a 12-hour light/dark cycle. Animals were fed a standard diet (A04, SAFE Diet, Augy, France). All procedures were conducted in accordance with institutional animal care guidelines and were approved by the Institutional Ethics Committee No. 16, Paris, France (license number 16-090). Throughout the study, mice underwent regular assessments including body weight monitoring and metabolic evaluations. Mice were euthanized at over 18 months of age, regardless of the presence or absence of tumor-related signs. Liver tissues were collected and processed for subsequent analyses.

### Histologic characterization of tumors

Freshly collected mouse liver tissues were fixed in 10% neutral buffered formalin for 24–48 hours (fixative:tissue volume ratio of 10:1), transferred to 70% ethanol for 24 hours before paraffin embedding. Paraffin blocks were cut into 5-µm-thick sections, which were subsequently dried overnight at 37 °C and stained with Haematoxylin-Eosin-Saffron (HES) using an automated slide stainer before digitalization. The description and diagnosis of neoplastic and non-neoplastic mouse liver lesions followed the International Harmonization of Nomenclature and Diagnostic Criteria (INHAND) guidelines for Lesions of the Rat and Mouse Hepatobiliary System and the accepted consensual terminology used in mouse pathology, as published by international committees and experts (Thoolen *et al*, 2010; Deschl *et al*, 2001). The researcher performing the histological evaluation (P.C.) was not blinded to sample identity at the time of the analysis. Liver neoplasms were evaluated for tumor architecture and relationship with adjacent tissue, growth pattern, areas of necrosis, neoplastic cell morphology, cytonuclear atypia, stromal composition, and leukocyte infiltration. Regional or distant metastasis, invasive growth or lympho-vascular invasion, tumor size, high mitotic activity, and severe atypia were considered key criteria of malignancy by decreasing order of importance.

### Whole exome sequencing and copy number alterations

Samples were sequenced to an average depth of 201× and 194× for the tumor and normal samples, respectively. Raw reads were aligned to the mouse reference genome (UCSC mm10) using BWA-MEM. Then, duplicated reads were marked with Picard Tools, and base quality scores were recalibrated using GATK (Van der Auwera *et al*, 2013). Somatic variant calling was performed using 12 different variant callers FreeBayes (Garrison & Marth, 2012), LoFreq, MuSE (Fan *et al*, 2016), Mutect (Cibulskis *et al*, 2013), Mutect2 (McKenna *et al*, 2010), Pindel (Ye *et al*, 2009), Scalpel (Fang *et al*, 2016), Seurat (Christoforides *et al*, 2013), SomaticSniper (Larson *et al*, 2012), Strelka (Kim *et al*, 2018), VarDict (Lai *et al*, 2016), and VarScan2 (Koboldt *et al*, 2012). A Panel of Normals (PON) was built using GATK Mutect2. An ensemble approach between the variant callers was used to filter false-positive variants. A minimum of five callers and four callers were required to retain a somatic SNV and indel, respectively. Finally, somatic variants found in the PON were filtered out.

Copy number alterations (CNA) were called using VarScan2 Copy Caller. Adjusted log2 ratio were then analyzed with circular binary segmentation as implemented in the DNA copy R/Bioconductor package (Olshen *et al*, 2004) to translate log2ratios measurements in regions of equal copy number. We used a threshold value of 0.1 to call the CNA alterations.

### Mutagenesis

Single base substitutions (SBSs) were described based on the pyrimidines of the Watson-Crick base pairs, resulting in six major substitution types (C>A, C>G, C>T, T>A, T>C, and T>G). These were further classified across their 16 trinucleotide contexts, defining 96 distinct SBS types.

### RNA Sequencing and Data Analysis

RNA sequencing was conducted by Novogene (Cambridge, UK). RNA integrity was assessed using the RNA Nano 6000 Assay Kit on a Bioanalyzer 2100 (Agilent Technologies, CA, USA). Total RNA was used for library construction. mRNA was isolated using poly-T oligo-attached magnetic beads and fragmented under elevated temperature in First Strand Synthesis Reaction Buffer. First-strand cDNA was synthesized using random hexamers and M-MuLV Reverse Transcriptase (RNase H-), followed by second-strand synthesis using DNA Polymerase I and RNase H. Overhangs were converted to blunt ends, 3’ ends were adenylated, and hairpin-loop adaptors were ligated. Fragments (370–420 bp) were purified using the AMPure XP system (Beckman Coulter), PCR-amplified with Phusion High-Fidelity DNA polymerase, and assessed for quality on the Bioanalyzer 2100. Indexed libraries were clustered using the cBot Cluster Generation System and TruSeq PE Cluster Kit v3-cBot-HS (Illumina), then sequenced on the Illumina NovaSeq platform to generate 150 bp paired-end reads.

Sequencing quality was assessed using FastQC ([http://www.bioinformatics.babraham.ac.uk/projects/fastqc/] (http://www.bioinformatics.babraham.ac.uk/projects/fastqc/)) and aggregated across samples with MultiQC (v1.7) (Ewels *et al*, 2016). Reads with a Phred quality score below 30 were filtered out to ensure high data quality.

High-quality reads were aligned to a customized mm10 mouse reference genome, including the mCherry transgene, using Subread-align (v1.6.4) (Liao *et al*, 2013) with default parameters. Gene-level counts were quantified with featureCounts (v1.6.4) (Liao *et al*, 2014).

Differential expression analysis was performed using DESeq2 (v1.26.0) (Love *et al*, 2014). To account for technical variability, a batch correction was applied based on sequencing run times. Genes with Benjamini–Hochberg adjusted P < 0.05 were considered significantly differentially expressed.

Principal Component Analysis (PCA) was performed using the prcomp function from R’s base ’stats’ package on variance-stabilized transformed (VST) counts. PCA allowed visualization of sample clustering and assessment of batch effects, ensuring biological replicates grouped together and that batch correction successfully minimized technical variation. GO enrichment analysis was conducted with clusterProfiler (v4.6.0) (Yu *et al*, 2012), focusing on Biological Process (BP) and KEGG pathway categories. A padj ≤ 0.05 was used as the significance threshold, considering an appropriate gene background. Heatmaps were generated using the heatmap.2 function (v3.1.3.1), based on DESeq2-normalized expression values. Venn diagrams were produced using Python’s matplotlib venn library. Custom R scripts were used to visualize gene signatures across DESeq2-normalized counts.

Ctnnb1 activated and axin1Δ signatures were sourced from Abitbol et al (2018). The Nrf2 signature was from (Mitsuishi *et al*, 2012a).

### Imaging Mass Cytometry

Imaging Mass Cytometry (IMC) was performed using the Hyperion™ Imaging System (Standard BioTools, USA), a multiplexed spatial proteomics platform that enables simultaneous detection of multiple protein markers at subcellular resolution in formalin-fixed paraffin-embedded (FFPE) tissue sections. IMC combines laser ablation with cytometry by time-of-flight (CyTOF) to detect metal-conjugated antibodies, overcoming limitations associated with fluorescence-based imaging such as spectral overlap and tissue autofluorescence (Giesen *et al*, 2014). Mouse liver tumor FFPE sections were processed according to standard protocols including dewaxing, rehydration, and heat-induced antigen retrieval. Sections were stained with a validated panel of 10 metal-tagged antibodies (AMK Biotech) following the steps previously described (Elaldi *et al*, 2021). Nuclear staining was performed using iridium intercalator. Regions of interest (ROIs) of approximately 1 to 2 mm² were selected on the tissue based on HES-stained sections to ensure representative sampling of both tumor and surrounding liver parenchyma and to avoid necrotic or hemorrhagic areas. Scale bars (200 µm) of the uncropped images are shown in Fig. EV4.

Image acquisition was conducted at a spatial resolution of 1 µm using the Hyperion Imaging System. Raw data were exported as OME-TIFF files via the MCD™ Viewer software (Standard BioTools, USA). For each marker of the used IMC-panel noise and background removal was performed using CellProfiler Software to generate an automatic binarization mask used to automatically remove the noise based on a learning-approach of pixel classification. All cleaned images were subsequently reviewed by an expert to ensure the absence of residual noise.

### Image analysis and quantification

Selected ROIs from denoised images of mice liver tumor sections were analyzed as follows: nuclei were segmented from the DNA1 channel using the legacy Cyto2 model, running on Cellpose 4.0.6 via the napari-serialcellpose plugin (v0.2.2) (Stringer *et al*, 2021). Cellprofiler 4.2.8 was used for subsequent quantification of immune marker positivity and relative areas versus total DNA. Metadata were extracted from file names. Integrated intensities of Cellpose-segmented nuclei were calculated for each marker. Since denoising reduced background fluorescence to near zero, any cell exhibiting signal above background was considered positive. The percentage of marker-positive cells relative to total cells (cellpose objects) was calculated per ROI. For area-based measurement, the areas of marker signals above the same threshold were compared relative to total DNA area per ROI. Data means were displayed as bar plots, with the individual ROI means shown as dots. Error bars represent standard error of the mean (s.e.m.). Representative images of segmented ROIs are displayed in figure panels for transparency. Cellprofiler pipelines are available upon request.

### Ki67 and gH2AX

Fresh liver was fixed in 10% phosphate-buffered formalin overnight. Paraffin wax sections of 5 µm were prepared for immunostaining (Ki67-abcam ab15580, gH2AX-Merck Millipore 05-636).

## Data availability

RNA-seq data are currently being deposited in the Gene Expression Omnibus (GSE).

## Acknowledgements

We thank Jean-Charles Graziano from the animal facility, the CRCM Integrative Bioinformatics (Cibi), preclinical TrGET platform, and the Experimental Histology platform (ICEP) for their support and Douglas Maya (IBIS, Sevilla) for helpful discussions. We gratefully acknowledge Frédéric Fiore (CIPHE, Marseille) for his guidance on generating mouse models and Marielle Bréau and Serge Adnot in the initiation of this project. This work has been supported by “La Ligue Nationale Contre le Cancer”, Equipe Labellisée, the “Institut National du Cancer” (INCA, PLBIO 2019), the Agence Nationale de la Recherche (ANR) (Grant THALATEL), the cross-cutting Inserm program on aging (AGEMED), the Inserm INTERAGING Program and the National Nature Science Foundation of China (32470830, 92478135).

## Author Contributions

L.B. and V.G. coordinated the study in collaboration with C.D. L.B. supervised tumor detection and sampling, prepared mRNA and genomic DNA for WES and RNA-seq analyses with assistance from M.B., and oversaw the generation of paraffin-embedded slides for histology and imaging. J.V. provided all RNA-seq data, while A.G. and F.B. performed WES analyses. P.C. and C.D. conducted the histological characterization of tumors. C.G. and A.M. performed Imaging Mass Cytometry, and T.E. conducted image analysis and quantification in collaboration with C.L. D.C. performed genomic analyses. L.G. performed database analyses, C.D., F.B., C.L and V. G. contributed to study funding. V.G. wrote the manuscript with input from all authors.

