## Supplemental data for "TERT drives liver tumorigenesis beyond telomere elongation"

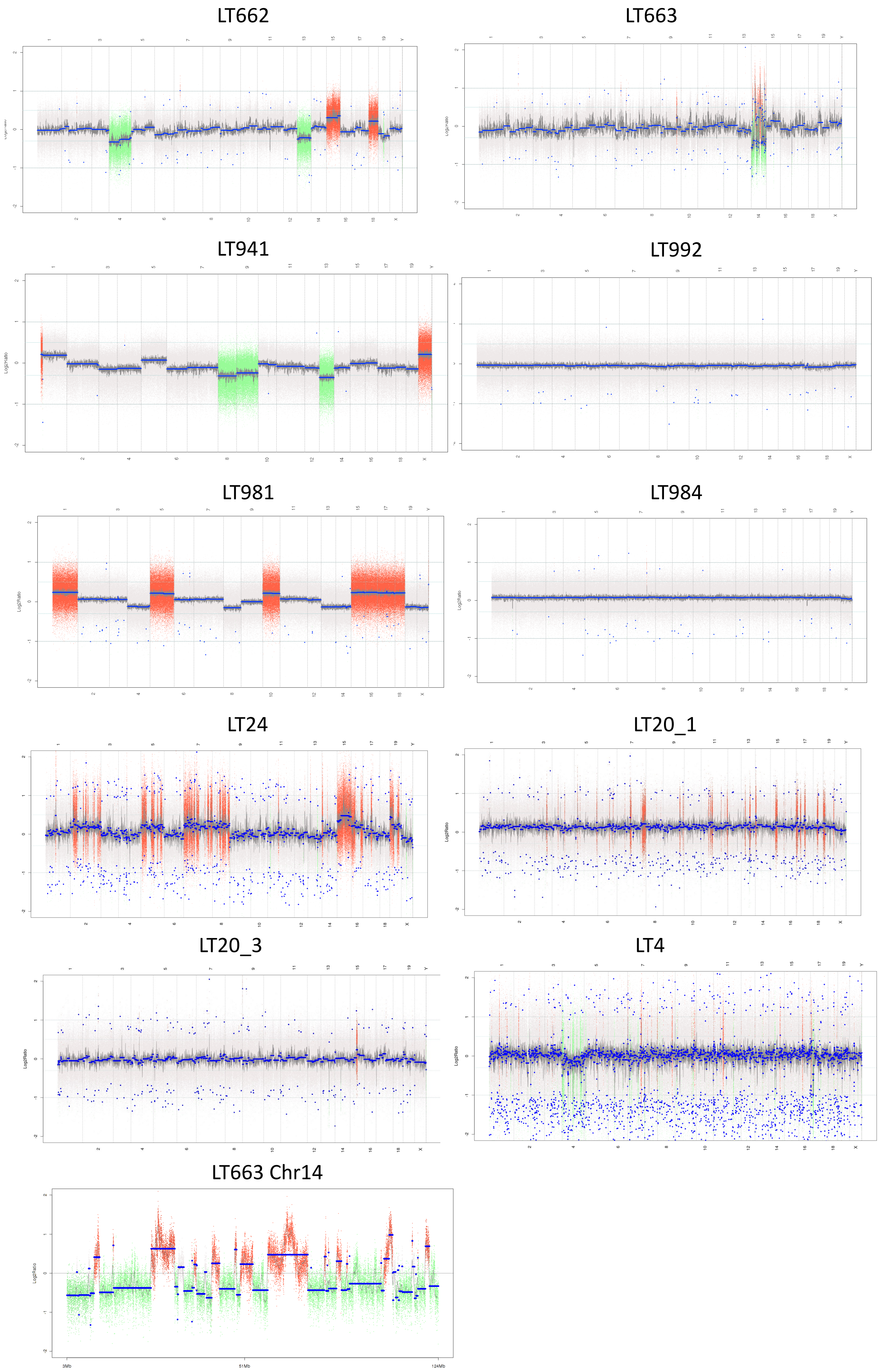


**Figure EV1. Copy-number-alteration (CNA) profiles in p21$^{+/Tert}$ and p21$^{+/TertCi}$ liver tumors.**

Gains and losses are indicated in red and green respectively. Tumors are indicated in the figure and chromosomes are numbered for each tumor.

Chymotrypsis on chromosome 14 of the LT663 tumor is shown at the bottom of the figure.

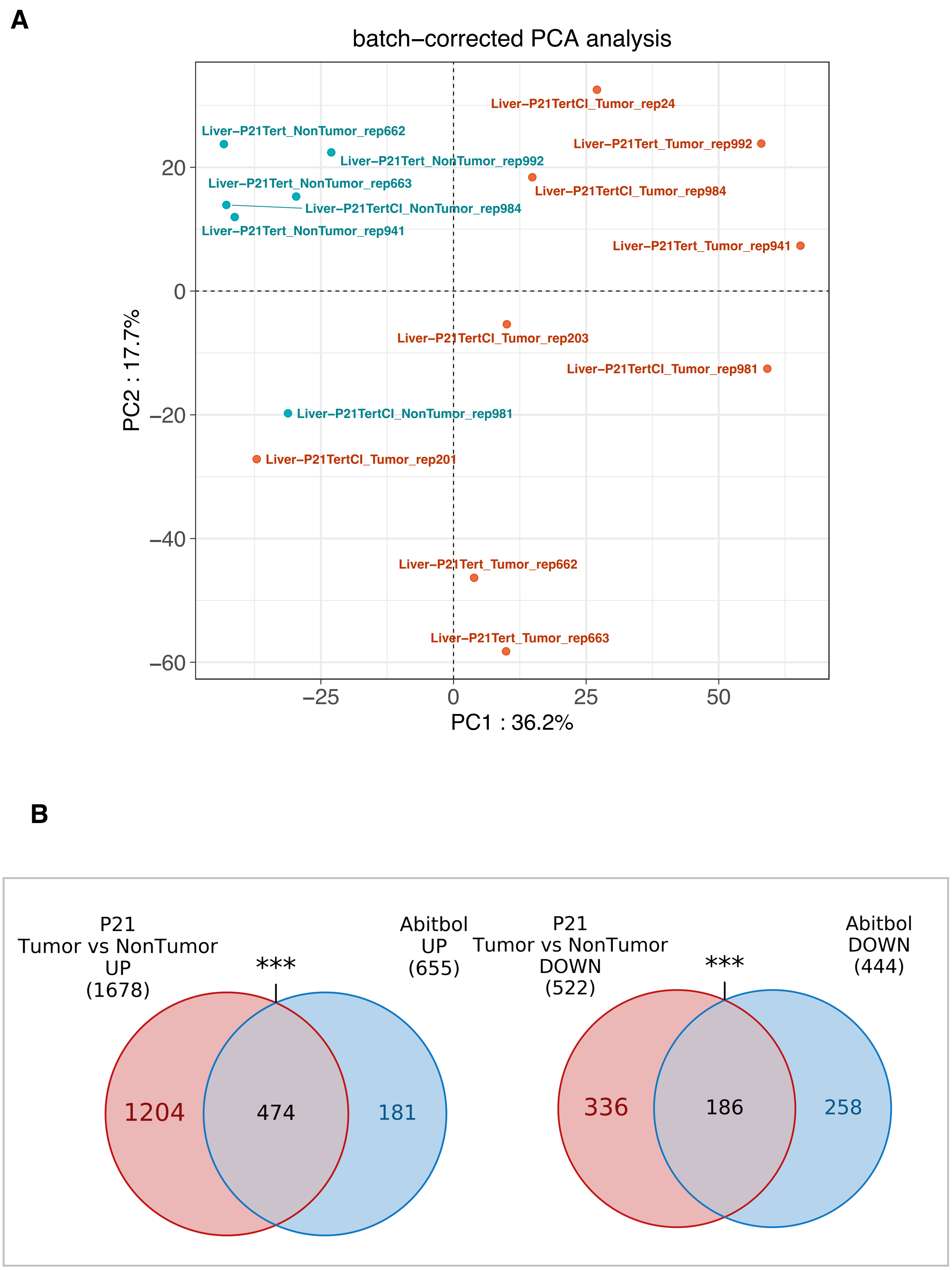


**Figure EV2. Transcriptomic profiles of $p21^{+/Tert}$ /$p21^{+/TertCi}$ tumors**

**A.** Principal component analysis (PCA) of tumor transcriptomes. PCA plots showing the clustering of liver tumors based on RNA-seq gene expression profiles**.** Each point represents an individual tumor sample, highlighting similarities and differences in global transcriptional patterns.

**B.** Comparison of transcriptomic profiles between $p21^{+/Tert}$ /$p21^{+/TertCi}$ tumors and *Axin1*Δ HCC. Venn diagram showing the number and overlap of significantly ($p<0.05$) upregulated genes (left panel) or downregulated genes (right panel) between $p21^{+/Tert}$ /$p21^{+/TertCi}$ tumors and HCC carrying an Axin1 deletion [48].

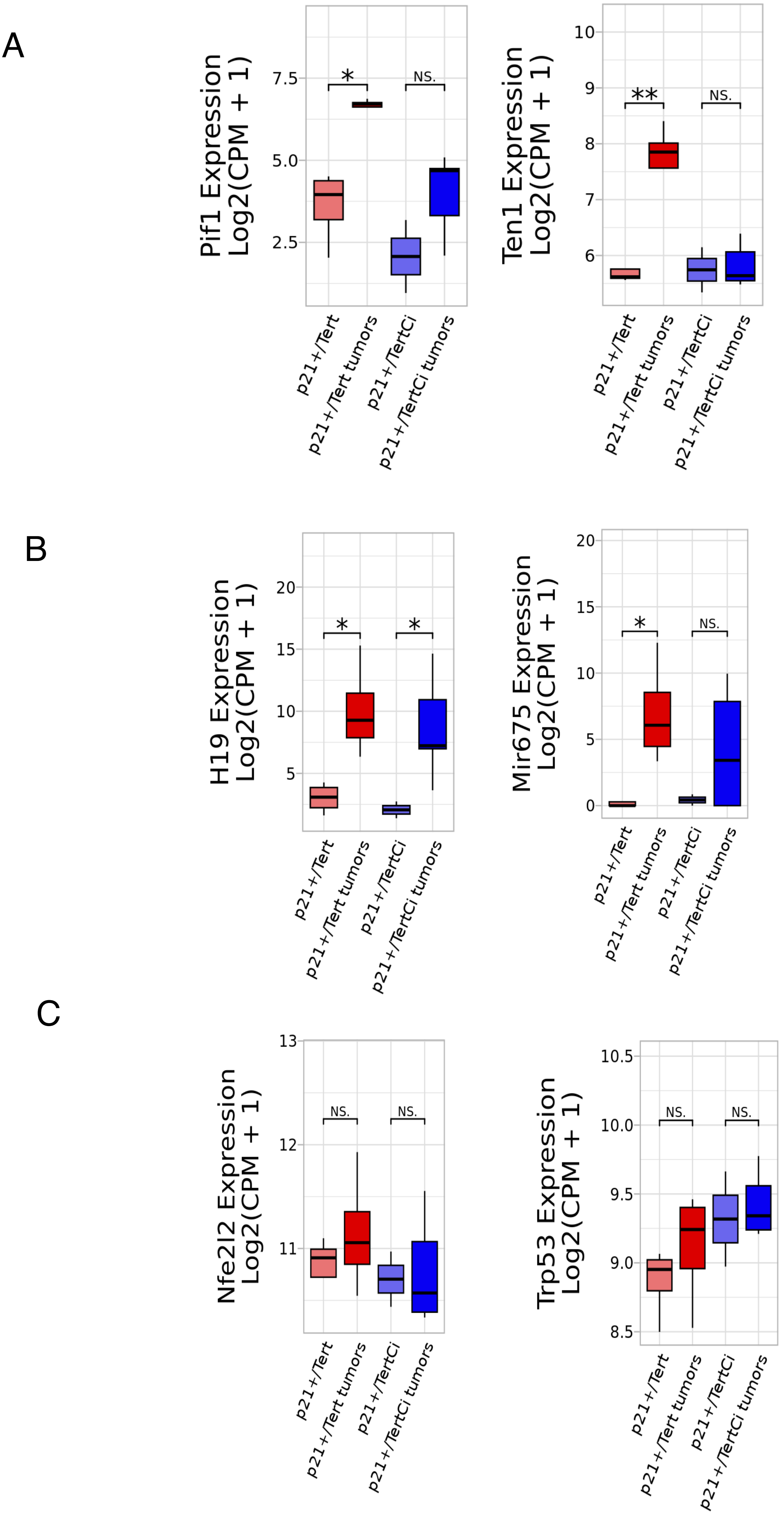


**Figure EV3. Expression levels in p21$^{+/Tert}$ and p21$^{+/TertCi}$ tumors of :**
A. Pif1 and Ten1,
B. H19 and Mir675,
C. Nfe2l2 and Tpr53.
Statistics *p<0.05, **p<0.01 vs non tumoral p21$^{+/Tert}$ or p21$^{+/TertCi}$

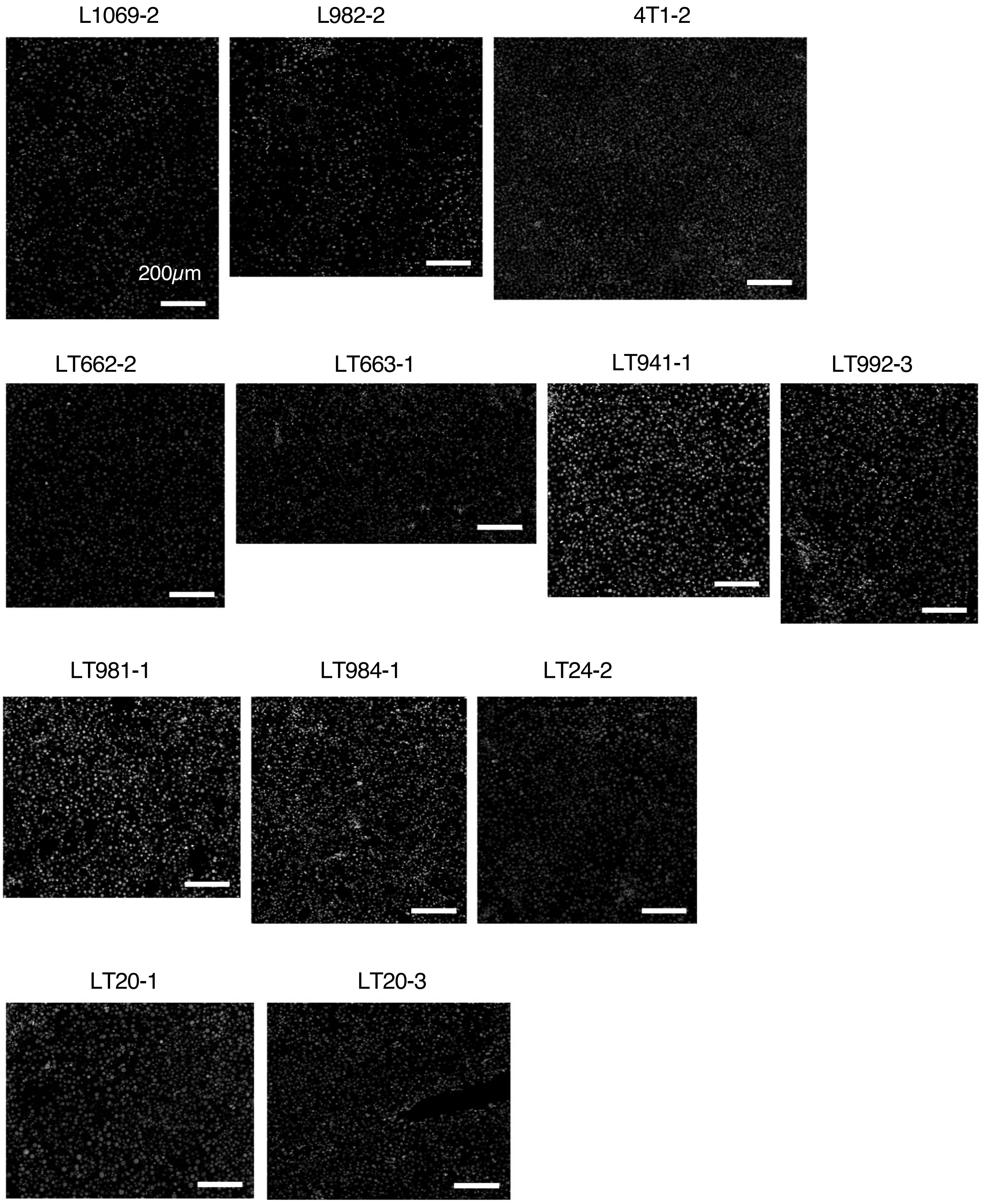


**Figure EV4. Scale bars for the uncropped images of p21$^{+/Tert}$ and p21$^{+/TertCi}$ liver and of the tumors**

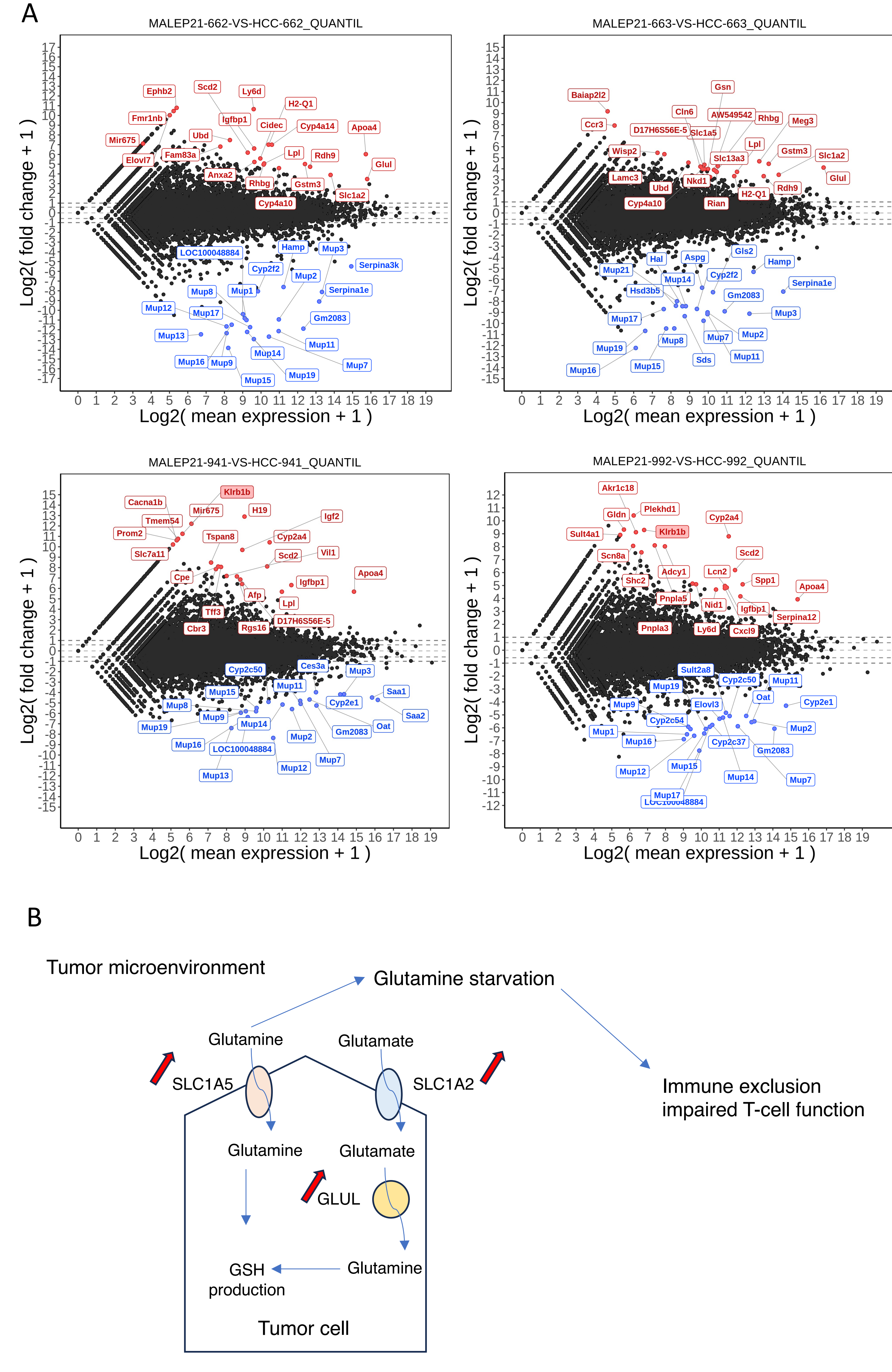


**Figure EV5. Expression changes in p21$^{+/Tert}$ tumors.**

**A.** The MA-plots (Mean–Average plot) highlights the genes that are most differentially expressed in the indicated tumors.

**B.** Proposed model explaining tumor immune evasion in LT662 and LT663. Both beta-catenin activated tumors establish a glutamine-depleted microenvironment consistent with the immune-excluded phenotype. Increased glutathione (GSH) production may also contribute to enhance antioxidant defences.
